# Wnt/β-catenin signaling regulates fibrotic atrophy of intra-articular adipose tissue in post-traumatic osteoarthritis

**DOI:** 10.64898/2026.06.08.730865

**Authors:** Alexander J. Knights, Danny M. Nguyen, Seth Kahan, Michael D. Newton, Huong X. Tran, Aanya Mohan, Isabelle J. Smith, Neil Bhate, Lindsey Lammlin, Stephen J. Redding, Luke Stasikelis, Tong Yang, Rida Pervez, Madison Buckles, Erica L. Scheller, Kurt D. Hankenson, Tristan Maerz

## Abstract

Synovial joints like the knee are home to adipose tissue depots whose anatomy and functions are closely intertwined with that of other intra-articular soft tissues such as synovium, underscoring the growing understanding that joints are multi-tissue organs. Traumatic joint injury and the onset of osteoarthritis (OA) dramatically remodel the intra-articular adipose niche, marked by infiltration of fibrotic tissue postulated to underpin OA-associated joint stiffness and pain, yet we know very little about the disease-associated dynamics of joint adipose remodeling nor the mechanisms driving these phenomena. Here, we employed 2D histomorphometry and spatial transcriptomics, alongside 3D osmium tetroxide-enhanced micro-computed tomography to comprehensively define the spatiotemporal, structural, and transcriptional rewiring of joint adipose tissue in a non-invasive mouse model of post-traumatic osteoarthritis (PTOA). These revealed marked loss of intra-articular adiposity accompanied by expansion of fibroblast-rich, collagen-dense tissue with pro-fibrotic hallmarks and Wnt/β-catenin-enriched gene programs. Joint adipose exhibited a distinct transcriptional signature compared to subcutaneous white adipose tissue, pointing to unique, depot-specific functions. Stromal cells isolated from PTOA joints had heightened baseline expression of fibrotic and Wnt pathway genes and exhibited impaired *de novo* adipogenesis, in contrast to cells derived from healthy joints. In accordance with the destabilized biomechanics of PTOA joints, *in vitro* modeling demonstrated that prolonged, injurious loading and perturbed Wnt/β-catenin signaling were convergent anti-adipogenic cues that suppressed lipid droplet formation and adipogenic gene induction, while promoting markers of fibrosis in joint-derived stromal cells. Complementary gain-of-function studies using *ex vivo* joint adipose explants and *in vivo* joint injections demonstrated that chronic Wnt/β-catenin activation, as seen in OA joints, is sufficient to diminish the intra-articular adipogenic program and shift adipose to a more fibrotic phenotype, independent of joint injury. Collectively, these findings establish a multi-modal framework for quantifying joint adipose atrophy and implicate aberrant Wnt/β-catenin signaling and pathological mechanical loading as key factors impairing *de novo* adipogenesis and driving fibrotic remodeling of intra-articular adipose tissue in PTOA.

## Introduction

Interest in the contributions of systemic metabolism and adipose tissue to joint disease is surging. Recent work has highlighted adipose tissue as an important mechanistic regulator of osteoarthritis (OA) ^1,2^, a debilitating joint disease characterized by pain, fibrosis, inflammation, and cartilage loss. However, relatively little attention has been given to local, intra-articular adipose depots that exist in joints like the knee, and whose functions at steady-state and in disease are not well understood. Despite its high prevalence, there are no disease-modifying treatments for OA, and extensive additional research is needed to understand which specific tissues, cell types, and molecular processes drive disease onset and progression. Fibrosis of adipose tissue is a hallmark of various chronic diseases^3–6^, but fibrosis of intra-articular adipose tissue and the mechanisms that drive it are poorly studied to-date.

The infrapatellar ‘fat pad’ (IFP) is the largest intra-articular adipose depot, situated behind the kneecap and historically believed to largely function as a shock absorber in the knee joint^7,8^. The IFP develops from embryonic *Gdf5*-lineage cells^9–11^, and continues to be populated postnatally by adipocytes that arise from local stromal progenitors expressing *Dpp4*^12^, which have also been characterized in other adipose depots^13–15^. It forms an integrated functional and anatomic unit with synovium^12,16,17^, the connective tissue that lines synovial joints, and is structurally homologous in humans and rodents. Although adipose tissue is distributed throughout the knee joint, the size and unified structure of the IFP has made it the focal point of clinical studies aimed at understanding joint adipose tissue in OA. Interestingly, the clinical literature has shown no correlation between BMI and joint adiposity^8,18–21^, in contrast to most major adipose depots in the body. Yet, there are conflicting reports on the association between intra-articular adipose tissue and OA, with some studies positively correlating IFP size with disease status^21,22^, others showing the opposite^8,17,23–25^, and others finding no correlation^18,26^. What is clear, however, is that stiffness and inflammation of the IFP are associated with elevated pain and OA severity^23,24,26–28^, suggesting that both ‘quantity’ and ‘quality’ of intra-articular adipose tissue are worthy of investigation.

Experimental modeling of OA in a pre-clinical setting allows us to remove many of the confounding factors that limit clinical studies, such as metabolic status, age, lifestyle, and known (or unknown) history of trauma. Joint injury is a leading risk factor for developing OA, termed post-traumatic osteoarthritis (PTOA) ^29^, and various surgical and non-surgical pre-clinical models exist for inducing PTOA. Our group has extensively characterized the non-surgical anterior cruciate ligament rupture (ACLR) model, which faithfully recapitulates the clinical mechanism of ACL injury and leads to PTOA, including hallmark phenotypic manifestations such knee pain, joint dysfunction, inflammation, fibrosis, cartilage degeneration, and pathological mineralization^30–35^. Studies by our group and others have consistently demonstrated that overactive Wnt/β-catenin signaling accompanies these pathological changes in PTOA^36–42^, and Wnt is known to be anti-adipogenic and pro-fibrotic^43–47^, highlighting its potential relevance to joint adipose biology and pathology. Remodeling of the intra-articular adipose niche has been noted in the pre-clinical arthritis literature, with the Griffin group showing that the IFP diminishes with age in rats^48^, then later observing histological IFP atrophy in the ACLR mouse model^49^. This histological observation was corroborated in other rodent and rabbit studies of joint injury^50–52^. Other studies have shown evidence of pathological IFP remodeling acutely post-injury, prior to phenotypic pain sensitization^53,54^, suggesting that this highly neurovascular niche may drive pain in OA. An insightful study from McClure *et al* sought to dissect the contributions of joint adipose tissue from systemic depots, suggesting that intra-articular adipose may be protective against joint pathology under obesogenic conditions, in a sex-dependent manner^10^. Relevant to this, intra-articular adipose tissue does not assume a pathological inflammatory phenotype to the same extent seen in large peripheral adipose depots like visceral adipose tissue^19,55^, suggesting that intra-articular adipose itself may have distinct intrinsic functions and biology. Recent single-nuclei RNA-sequencing experiments performed on IFP tissue have enabled unprecedented insights into the transcriptional signatures and functions of the adipogenic lineage in the joint^56–58^, that were not previously afforded in the early wave of single-cell RNA-sequencing.

Here we sought to characterize the spatiotemporal dynamics and functions of intra-articular adipose tissue, under steady-state conditions and in PTOA. We shed light on conserved and distinct functions of intra-articular adipose tissue compared to subcutaneous adipose tissue; robustly evaluated the healthy joint-wide distribution of adipose tissue and demonstrated its loss in PTOA; and identified a pathological intersection between overactive Wnt/β-catenin signaling and mechanical loading that may underpin the observed adipose atrophy in the joint.

## Results

### Intra-articular adipose tissue undergoes progressive fibrotic atrophy during post-traumatic osteoarthritis

Fibrotic and inflammatory remodeling of intra-articular soft tissues, notably synovium, is well reported in studies of human and pre-clinical PTOA^31,59,60^. However, concurrent changes to joint adipose tissue, especially in the early phase after injury, have been less well characterized. In the non-invasive ACLR mouse model of PTOA, we found that genes closely associated with fibrosis were upregulated acutely in synovium and intra-articular adipose tissue following joint injury, and persisted as disease progressed (Fig 1A). On the other hand, adipogenic genes such as *Pparg*, *Plin1*, *Adipoq*, and *Fabp4* were strongly downregulated. This observation was corroborated in published human data^58^ that we re-analyzed, wherein fibrotic genes were elevated while adipogenic genes were decreased in synovial and IFP tissue from patients undergoing total knee arthroplasty compared to those without a history of arthritis (Fig 1B). To validate whether these gene signature shifts coincided with diminished intra-articular adipose tissue in PTOA, we performed 2D histomorphometry on Safranin O/Fast Green-stained sagittal knee sections from mice subjected to ACLR or their healthy (Sham) counterparts. In this model, 7d ACLR corresponds to the acute, inflammatory phase of disease, while 28d ACLR represents established PTOA^31,35^. In the anterior compartment, largely comprised of the IFP, the relative area of adipose coverage was significantly diminished by 28d post-ACLR, while in the posterior compartment, significant loss was detected by 7d post-ACLR (Fig 1C-D, Fig S1A-F). Interestingly, this approach did not detect statistically significant loss of relative adipocyte area in the anterior compartment for female mice at any disease timepoint.

**Fig 1.**
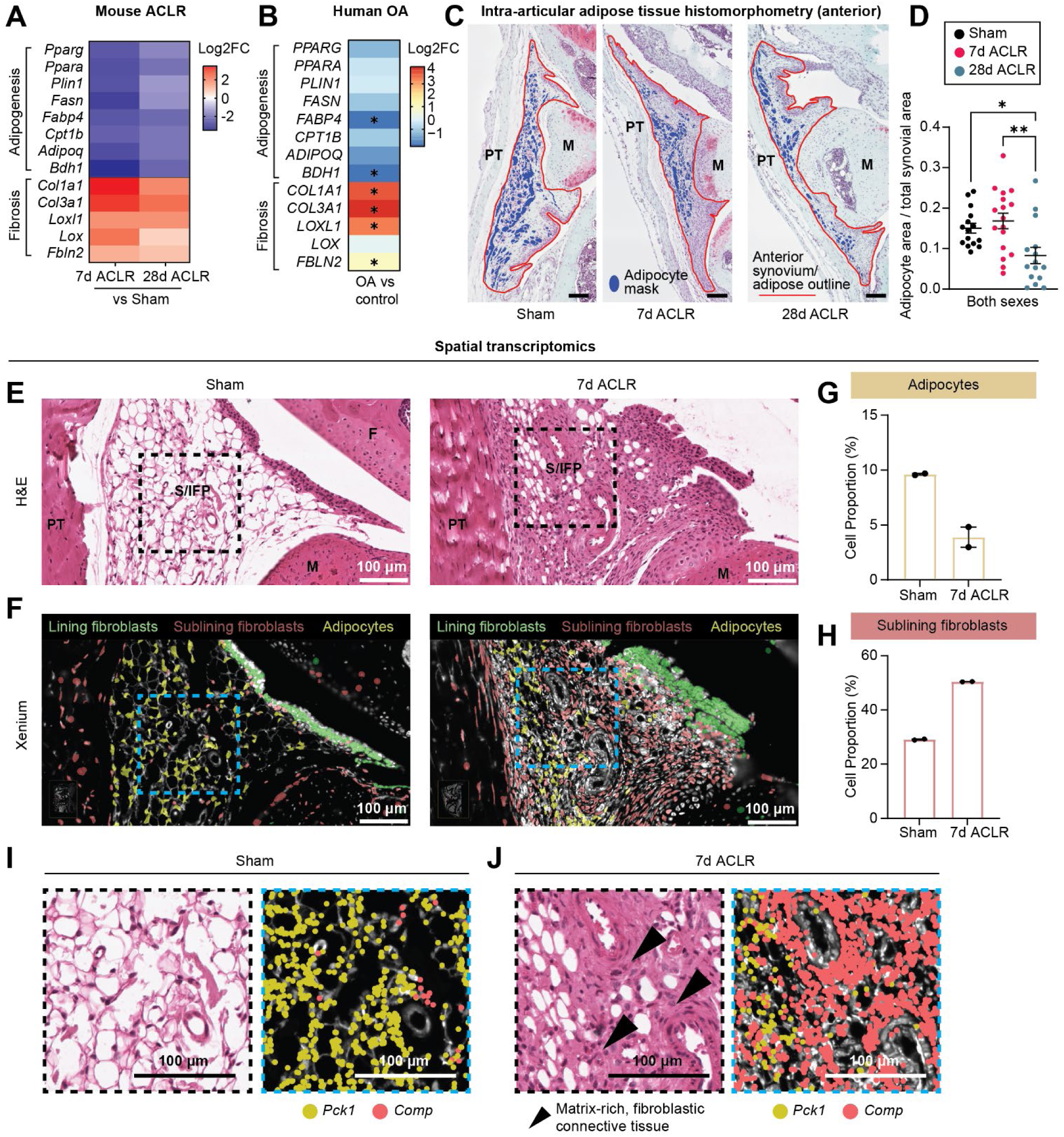
Intra-articular adipose tissue undergoes progressive fibrotic atrophy during post-traumatic osteoarthritis. (A) Bulk RNA-seq data of mouse synovium and intra-articular adipose tissue derived from GSE271903 showing relative expression (log2FC) of genes related to adipogenesis or fibrosis between 7d ACLR and Sham (healthy) or 28d ACLR and Sham. All genes are significant DEGs (*padj*<0.05). (B) Pseudo-bulk analysis of single-cell RNA-seq data of human IFP and synovium derived from GSE216651 showing relative expression (log2FC) of genes related to adipogenesis or fibrosis in OA tissues versus control tissue (without history of arthritis). Genes marked with an asterisk are significant DEGs (*padj*<0.05). (C-D) Representative images of anterior synovium and IFP from Sham, 7d ACLR, and 28d ACLR sagittal knee joint sections stained with Safranin O/Fast Green (C). Synovium/IFP is outlined in red; adipocyte mask is blue. (D) Ratio of anterior adipocyte area to total synovial outline area in Sham, 7d ACLR or 28d ACLR for both sexes (n=15-17 mice). Kruskal-Wallis testing with Dunn’s multiple comparisons test was performed where \**P*<0.05, \*\**P*<0.01. (E) H&E staining of sagittal knee joint sections from Sham (left) or 7d ACLR (right) male mice, focused on the anterior synovium and IFP. (F) Xenium spatial transcriptomics showing clustering of synovial lining fibroblasts, sublining fibroblasts, and adipocytes in the synovium/IFP from the same sections as in (E). (G-H) Proportion of adipocytes (G) and sublining fibroblasts (H) relative to all synovium/IFP cells in Sham and 7d ACLR Xenium sections (n=1 male and female mouse per condition). (I-J) Magnified fields of view from (E-F), demarcated by black (H&E) or blue (Xenium) dashed boxes. *Pck1* and *Comp* transcript expression is overlaid for both Sham (I) and 7d ACLR (J) Xenium views, with black arrowheads indicating emergent fibroblastic, matrix-rich tissue regions. ACLR: anterior cruciate ligament rupture. PT: patellar tendon. S/FP: synovium/IFP. M: meniscus. F: femur. Scale bars: 100 μm. Errors bars are means±SEM for (D, G-H).

To complement this histomorphometric approach and greatly increase cell identity resolution, we performed spatial transcriptomics on Sham and ACLR mouse knee joints using the 10x Xenium v1 platform (Fig S2A-C). Semi-supervised clustering and targeted gene signature profiling identified major intra-articular cell types, including lining and sublining fibroblasts, and adipocytes (Fig S2D-E). H&E post-staining of Xenium sections showed diminished adipose area alongside increased fibrotic matrix and cellularity in the anterior synovium and IFP at 7d post-ACLR (Fig 1E, Fig S2F). Overlay of major cell type clusters on the anterior synovium and IFP shed light on the cellular dynamics underpinning our histological observations; lining fibroblast accumulation corresponded to thickening of the lining layer, significantly increased sublining fibroblast cellularity was observed throughout ACLR tissue, and crucially, the relative proportion of adipocytes was diminished in ACLR compared to Sham joints (Fig 1F-H, Fig S2G). Close examination revealed that adipocytes, characterized by their large lipid droplets and expression of the metabolic enzyme *Pck1*, were partially replaced in ACLR tissue by emergent fibroblastic cells expressing the fibrotic matrix glycoprotein *Comp* (Fig 1I-J, Fig S2H-I). Importantly, these findings provide spatial corroboration of our prior characterization of emergent disease-associated fibroblast subsets in PTOA^37,61^; namely, progressive accumulation of synovial lining fibroblasts that likely underpins clinically-observed lining hyperplasia^62,63^, and dramatic emergence of sublining fibroblasts that leads to fibrosis^62,64^, driven by local proliferation. Together, histomorphometric analyses and spatial transcriptomics demonstrate the loss of intra-articular adipose tissue and adipocytes following joint injury and during PTOA progression.

### Osmium tetroxide-enhanced µCT reveals volumetric atrophy of intra-articular adipose tissue

Since adipose tissue is not uniformly distributed throughout the joint, and given the inherent limitations of 2D histomorphometry, we next sought to extend our findings by evaluating volumetric fat loss in the entire intra-articular space in a three-dimensional manner. Osmium tetroxide is a lipophilic heavy metal compound that has been previously used as a micro-computed tomography (µCT) contrast agent to visualize and quantify bone marrow adiposity^65^. However, this approach has never been harnessed for assessing joint adiposity. After fixing freshly isolated 28d ACLR and contralateral (uninjured) hindlimbs, we incubated them in 1% osmium tetroxide for 48 h, prior to high-resolution µCT scanning and quantitative analysis. 3D visualization reinforced that intra-articular adipose tissue is concentrated in the anterior compartment (primarily the IFP) and posterior compartment. Limited but nonetheless detectable deposits of adipose tissue were observed in the intercondylar space (i.e. pericruciate fat) and medial and lateral compartments, unlike synovium which envelops the entire joint space (Fig 2A-B). This distribution is consistent with our understanding of adipocytes - concentration into organized adipose depots as well as dispersed accumulation within and around parenchymal tissues^66^. Volumetric analysis supported our histomorphometric findings, showing significantly reduced adipose tissue in 28d ACLR joints across both compartments (34% reduction in anterior; 79% reduction in posterior) (Fig 2C-D).

**Fig 2.**
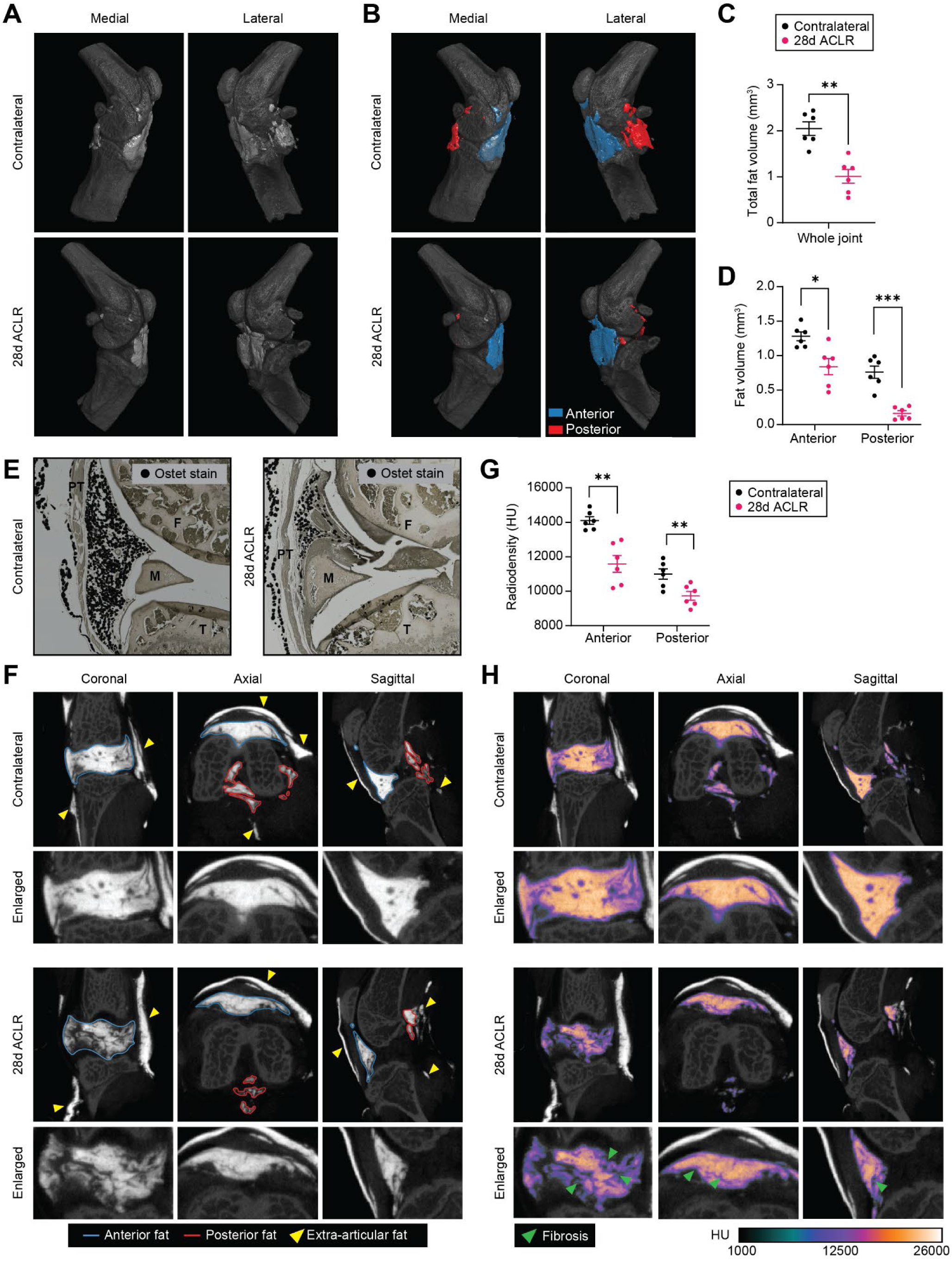
Osmium tetroxide-enhanced µCT reveals volumetric atrophy of intra-articular adipose tissue. (A-B) Representative 3D µCT renderings of contralateral and 28d ACLR knees stained with osmium tetroxide (Ostet) showing intra-articular adipose in grayscale (A) and colored by anterior (blue) or posterior (red) compartment (B) (n=6 mice). (C-D) Total (C) and compartmental (D) volume of intra-articular adipose tissue. (E) Sagittal histological sections stained with Ostet (black) contralateral and 28d ACLR joints. (F) Representative 2D µCT views of Ostet-stained intra-articular adipose in grayscale and outlined according to anterior (blue) or posterior (blue) compartment, with enlarged views below. Yellow arrowheads denote extra-articular adipose tissue regions not included in analysis. (G) Radiodensity of Ostet signal measured in Hounsfield Units (HU) in the anterior or posterior adipose compartment. (H) Representative 2D views with radiodensity colormapping overlaid (HU), including enlarged views. Green arrowheads denote fibrotic regions. For (C-D, G), two-tailed paired t-tests were performed to compare laterality, where \**P*<0.05, \*\**P*<0.01, \*\*\**P*<0.001. PT: patellar tendon. T: tibia. M: meniscus. F: femur.

Histological examination of osmium tetroxide-stained limbs verified complete tissue penetration and selective staining of lipid within adipocytes (Fig 2E). Extra-articular adipose tissue depots, such as subcutaneous adipose tissue around the joint, were visible by µCT but spatially distinct and readily excluded from intra-articular analysis (Fig 2F). In addition to reduced total adipose volume, tomographic radiodensity of joint adipose tissue was also diminished in PTOA joints, indicative of decreased contrast agent uptake due to reduced lipid concentration, and the pattern of signal loss observed in PTOA joints corresponds to the emergence of fibrotic tissue (Fig 2G-H). Overall, we developed a novel approach for 3D volumetric analysis of joint adipose tissue that demonstrates compartment-specific adipose loss in PTOA, corroborating and extending our 2D analyses.

### Intra-articular adipose is transcriptionally distinct from classical white adipose tissue

Intra-articular adipose tissue is proposed to have unique functions and composition^7^, distinct from classical white adipose tissue (WAT) depots of the body. These include physical insulation, shock absorption, lubrication, and serving as a putative reservoir for regenerative cells in the joint. Anatomically, intra-articular adipose tissue is integrated with synovium, and pathological remodeling is seen at the interface of the synovial sublining and adipose tissue during OA. To characterize the transcriptional signature of intra-articular adipose tissue, we performed bulk RNA-sequencing (RNA-seq) of the IFP and inguinal subcutaneous WAT of healthy adult mice (Fig S3A). Despite their common identity as adipose tissues, principal component analysis revealed distinct clustering of the tissues based on gene signature (Fig 3A). There were 5,753 significantly differentially expressed genes (DEGs, *padj*<0.05, log2FC>|0.585|, FPKM>1) between the two tissues (Fig 3B, Table S1). These DEGs underpinned enrichment of biological pathways and functions across a range of domains (Fig 3C-D, Fig S3B, Table S2). The IFP was enriched for musculoskeletal-related gene ontology terms including extracellular matrix assembly, elastic fiber formation, hyaluronan metabolism, chondrocyte differentiation, bone mineralization, and ossification.

**Fig 3.**
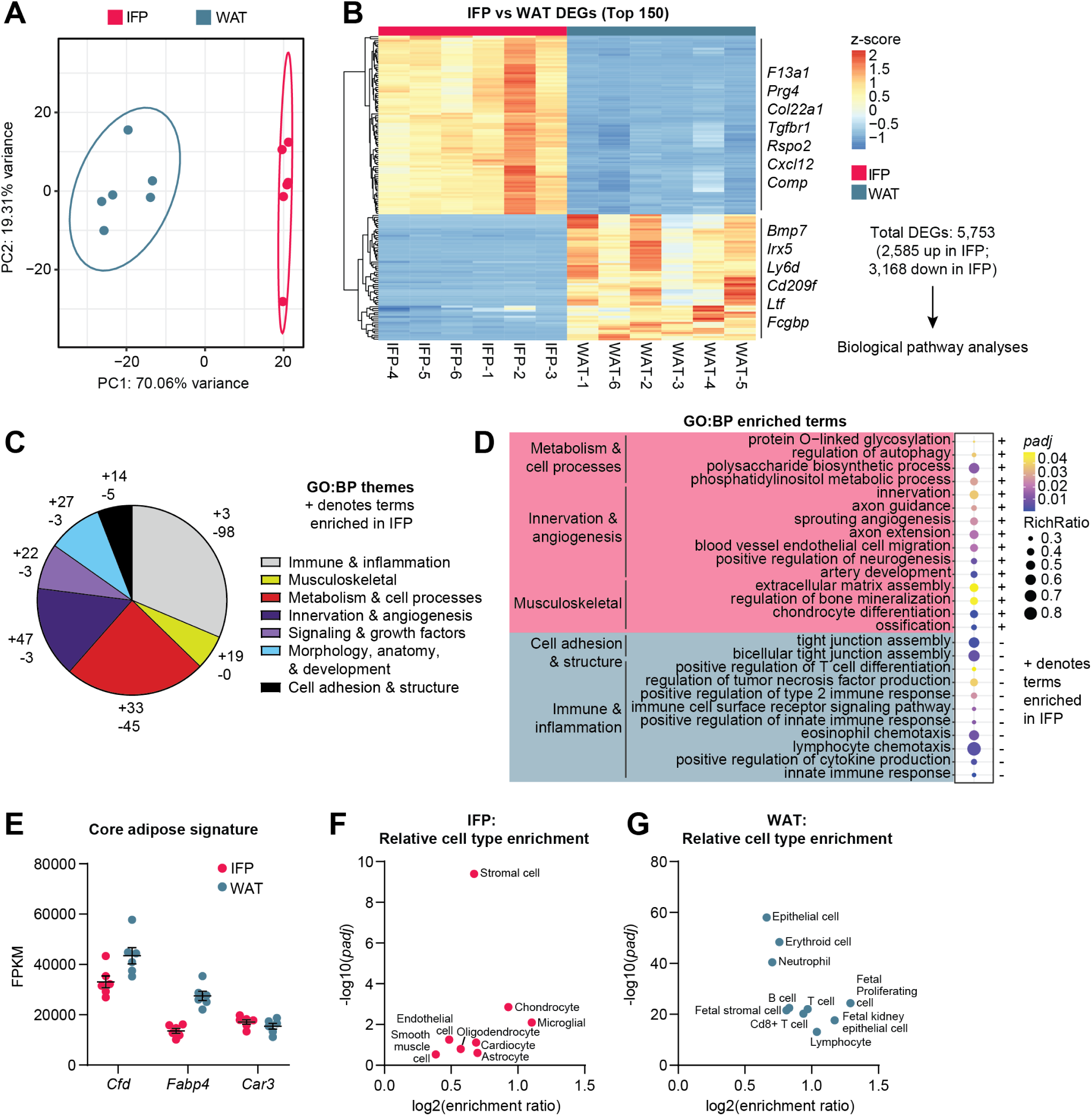
Intra-articular adipose is transcriptionally distinct from classical white adipose tissue. Bulk RNA-seq was performed on IFP and inguinal subcutaneous white adipose tissue (WAT) (n=6 mice). (A) Principal component analysis showing sample clustering based on tissue type. Ellipses represent the 95% confidence interval. (B) Hierarchical clustering of relative expression (z-score) of the top 150 differentially expressed genes (DEGs) between IFP and WAT (*padj*<0.05 and log2FC>|0.585|), with key DEGs noted. Total number of DEGs, up and down, are shown. (C) DEGs from (B) were used as input for Gene Ontology: Biological Processes (GO:BP) statistical enrichment testing; shown is a breakdown of broad gene ontology themes, with (+) denoting the number of terms enriched in IFP compared to WAT, and vice versa for (-). (D) Enriched GO:BP terms and their tissue-type directionality (+ or -). Bubble color represents significance (*padj*) and bubble size indicates RichRatio (DEGs present in the term divided by the total genes in the term). (E-F) Cell type enrichment analysis using WebGestalt 2024 showing cell types predicted to be relatively enriched in IFP (E) or WAT (F) based on differential gene signatures between the two tissues.

Despite both tissues sharing a core adipose gene signature (Fig 3E), these findings are indicative of greater relative stromal cell content in the IFP, which was corroborated by cell type enrichment analysis (Fig 3F-G). The IFP is known to contain stromal progenitors with strong propensity for osteochondral differentiation^67,68^, and enrichment of these terms compared to WAT is not surprising given the proposed regenerative function that synovium and IFP may play in joint homeostasis and healing. IFP was also enriched for pathways pertaining to innervation, neurotrophic signaling, and angiogenesis, in line with its highly vascular anatomy, its high degree of innervation, and its strong link to joint pain. Enrichment of these pathways in the IFP was supported by overrepresentation of endothelial cell, oligodendrocyte, and astrocyte signatures (the latter two likely corresponding to peripheral glial cells) (Fig 3F). On the other hand, WAT was comparatively enriched for adhesion, immune and inflammatory terms like chemotaxis, cytokine production, and tight junction assembly. Cell type enrichment analysis further confirmed greater comparative immune cell content in WAT, namely neutrophils and lymphocytes (Fig 3G). In summary, transcriptional profiling of IFP and WAT highlighted their distinct gene signatures, revealing greater transcriptional activity related to innervation, angiogenesis, and extracellular matrix remodeling in the IFP compared to WAT.

### Impaired adipogenesis in stromal cells from osteoarthritic joints

Given the unique gene signature and functions of intra-articular adipose tissue that we inferred, we next evaluated the adipogenic capacity of stromal cells derived from the IFP, and whether cells from PTOA IFP have compromised adipogenic differentiation. First, inguinal WAT was digested to yield the stromal vascular fraction, and these cells were cultured and subjected to adipogenic differentiation as a reference point (Fig 4A). WAT-derived stromal cells kept in growth medium retained a fibroblastic morphology and did not develop lipid droplets, whereas cells in the presence of adipogenic medium exhibited robust lipid droplet formation, induction of adipogenic genes, and strongly elevated nuclear expression of PPARɣ protein (Fig 4B-C, Fig S4D). Stromal cells derived from digested IFP of healthy mice were subjected to the same adipogenic differentiation and showed comparable lipid droplet formation, adipogenic gene induction, and nuclear PPARɣ accumulation, with no appreciable differentiation in the presence of growth medium (Fig 4B-C, Fig S4A-D). However, differentiation was significantly blunted in cells derived from PTOA IFP, which exhibited comparatively diminished lipid droplet formation and adipogenic gene induction (Fig 4B-C).

**Fig 4.**
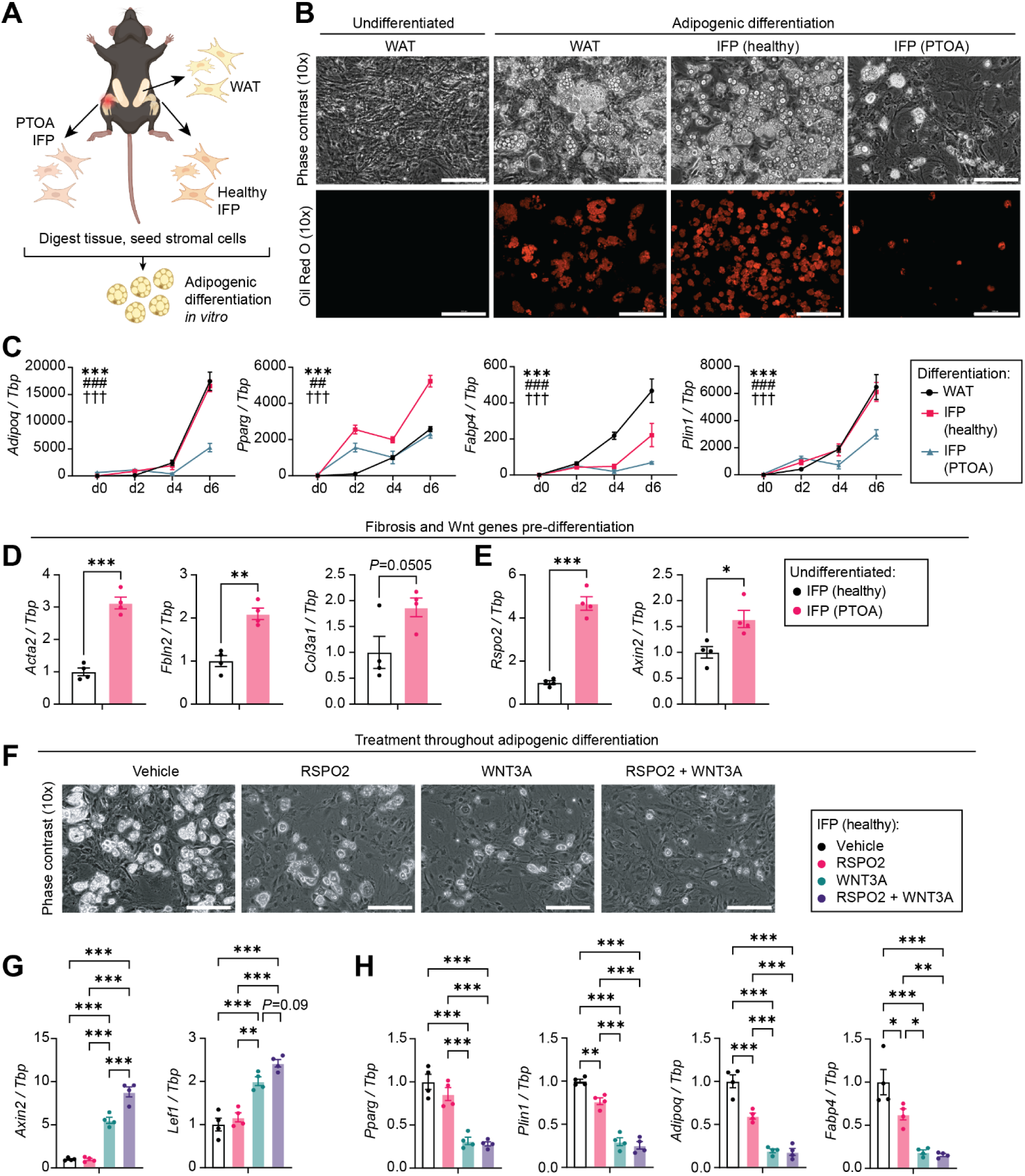
Impaired adipogenesis in stromal cells from injured joints. (A) White adipose tissue (WAT) and IFP/synovial tissue (healthy; or PTOA, 7d ACLR) were harvested from mice, tissue was digested, then stromal cells were seeded for culture prior to adipogenic differentiation. (B) Representative phase contrast (10x, top) and Oil Red O (10x, bottom) images showing morphology and lipid droplet formation in WAT-derived or IFP-derived stromal cells subjected to 6 days of adipogenic differentiation. Undifferentiated (growth medium only) WAT cells are shown on the left. (C) Relative expression of adipogenic genes over 6 days of differentiation for stromal cells derived from WAT, healthy IFP, or PTOA IFP (n=4). WAT d0 expression is set to 1 for each gene. Two-way ANOVA with Tukey’s multiple comparisons testing was performed. Significance is denoted by * (Interaction), # (Tissue), or † (Time), where one symbol indicates *P*<0.05, two symbols indicates *P*<0.01, three symbols indicates *P*<0.001. (D-E) Expression of fibrosis (D) and Wnt signaling (E) genes in undifferentiated stromal cells derived from healthy or PTOA IFP (n=4). Expression in healthy tissue was set to 1 for each gene. For (D-E), unpaired two-tailed t-tests were performed, where \**P*<0.05, \*\**P*<0.01, \*\*\**P*<0.001. (F-H) Healthy IFP stromal cells were treated with vehicle, RSPO2 (200 ng/mL), WNT3A (20 ng/mL), or RSPO2 + WNT3A throughout adipogenic differentiation. Representative 10x phase contrast images show lipid droplet accumulation (F). Expression of genes involved in Wnt signaling (G) or adipogenesis (H) (n=4). For (G-H), one-way ANOVA with Tukey’s multiple comparisons testing was performed, where \**P*<0.05, \*\**P*<0.01, \*\*\**P*<0.001. For all gene expression data (C-E, G-H), transcript levels were normalized to *Tbp* housekeeper expression, and error bars are means±SEM. Scale bars: 200 μm.

Prior to differentiation, PTOA IFP-derived cells had elevated levels of fibrotic genes *Acta2* (encoding αSMA), *Fbln2*, and *Col3a1*, compared to cells from healthy tissue, suggesting a less adipogenic and more fibrogenic predisposition (Fig 4D). The Wnt signaling pathway agonist *Rspo2* (encoding R-spondin 2/RSPO2) and the Wnt target gene *Axin2* were both elevated in stromal cells derived from PTOA IFP (Fig 4E). Wnt/β-catenin signaling is known to be anti-adipogenic and pro-fibrotic^43,69,70^, and treatment of healthy IFP-derived stromal cells during adipogenic differentiation with the potent Wnt/β-catenin activator CHIR99021 completely abrogated adipogenic gene induction and lipid droplet formation (Fig S4E-G), as was shown previously in adipogenic progenitors from human subcutaneous adipose tissue^70^. Wnt/β-catenin signaling is also known to be persistently overactivated in multiple joint tissues in OA, including synovium. We previously showed that R-spondin 2, a secreted protein responsible for amplifying Wnt/β-catenin signaling, is elevated in PTOA joints^37,39,71^ and has been reported to suppress differentiation of adipogenic progenitors^47^. Treating WAT-derived stromal cells with R-spondin 2 throughout adipogenic differentiation or at terminal differentiation blunted lipid droplet formation, and this was exacerbated with co-treatment with the Wnt ligand WNT3A (Fig S4H-I), consistent with the canonical role of R-spondin 2 in stabilizing Frizzled receptors that engage Wnt ligands. Similarly, treating healthy IFP-derived stromal cells with R-spondin 2 throughout adipogenic differentiation reduced adipogenic gene induction and lipid droplet formation, while synergism with WNT3A accentuated this effect and strongly increased expression of Wnt pathway genes *Axin2* and *Lef1* (Fig 4F-H). These results demonstrate that stromal cells from healthy IFP have comparable adipogenic capacity to WAT-derived cells, but cells from PTOA IFP have elevated fibrotic and Wnt gene expression at baseline and exhibit impaired adipogenesis *in vitro*. Overall, these findings suggest that the chronically overactive Wnt signaling observed in PTOA joints^38,39,42,72^ may limit *de novo* adipogenesis and contribute to diminished intra-articular adipose tissue in disease.

### Injurious mechanical loading disrupts adipogenic differentiation *in vitro*

Joint articulation naturally subjects intra-articular tissues to dynamic mechanical stresses, and these can become dysregulated and injurious in disease, such as in joint-destabilizing conditions like PTOA. Relatedly, disease-associated fibrotic remodeling, as observed in PTOA synovium and intra-articular adipose, alters tissue material properties that expose cells to persistent non-homeostatic loading conditions^73,74^. Compared to WAT, intra-articular adipose tissue has broader expression of mechanoreceptor genes including *Piezo1* and members of the transient receptor potential (*Trp*) family (Fig 5A). Notably, several of these mechanoreceptors are deregulated in human OA synovium and IFP compared to control tissue, and also show differential expression depending upon patient-reported pain status (Fig S5A-B).

**Fig 5.**
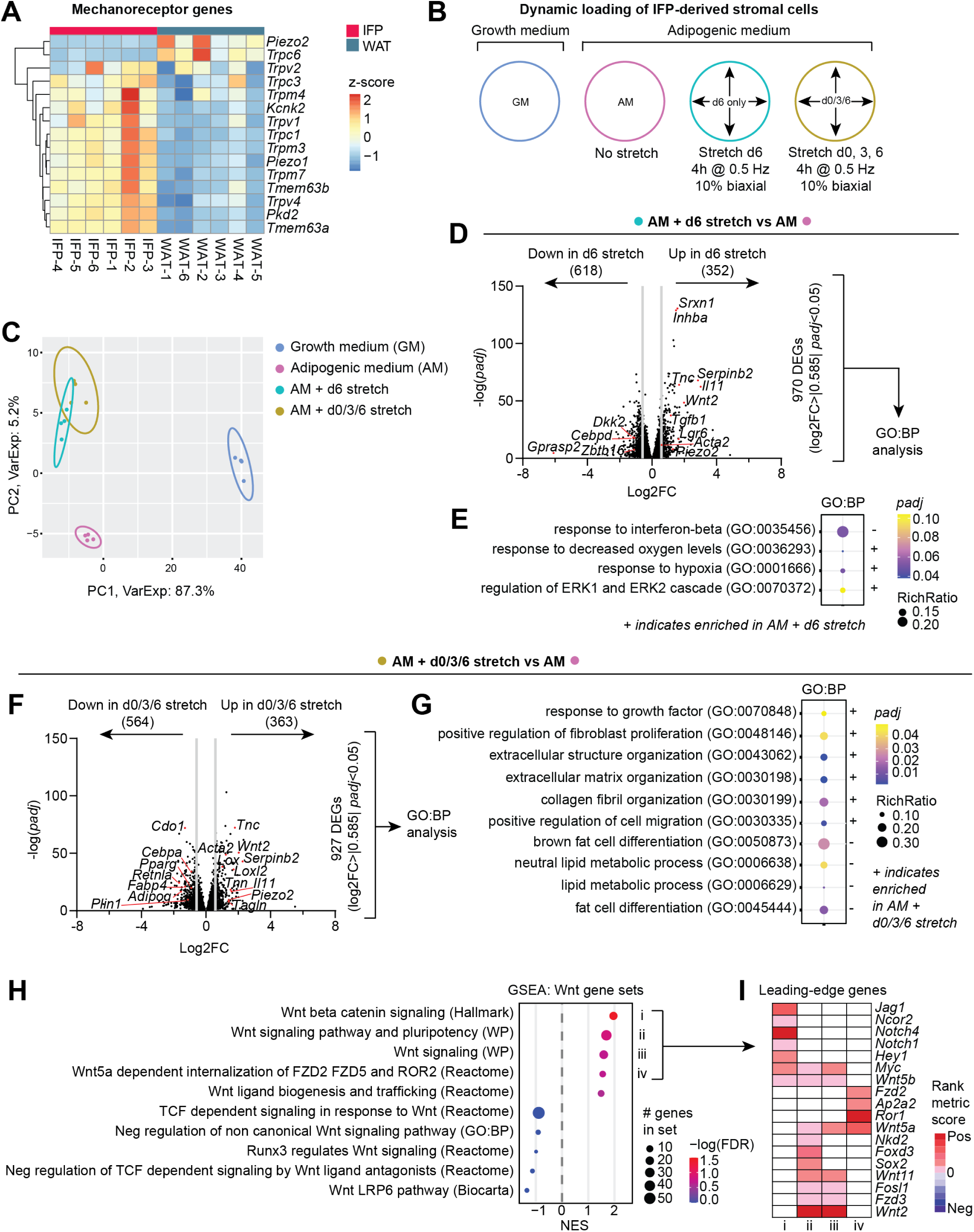
Injurious mechanical loading disrupts adipogenic differentiation *in vitro*. (A) Relative expression (z-score) of mechanoreceptor genes in healthy mouse IFP and subcutaneous WAT (n=6). Only genes with appreciable expression (FPKM>1) in either tissue are included. (B) Experimental design for FlexCell dynamic stretching. Healthy IFP-derived stromal cells were seeded onto 10 kPa BioFlex plates and grown in either growth medium (GM) or adipogenic medium (AM). Cells grown in AM were kept static throughout differentiation, or subjected to dynamic tensile stretching, either at terminal differentiation (AM + d6 stretch) or throughout differentiation (AM + d0/3/6 stretch). Loading bouts lasted 4 h, and were performed at 10% biaxial strain, 0.5 Hz. Cells were harvested 4 h following the final stretch (d6) or at the equivalent timepoint for static condition. (C) Principal component analysis (PCA) across RNA-seq conditions. Ellipses represent the 95% confidence interval. (D) Volcano plot showing significant DEGs between AM + d6 stretch versus AM alone (log2FC>|0.585|, *padj*<0.05). Key DEGs are named. (E) Pathway statistical enrichment analysis using the Gene Ontology: Biological Processes (GO:BP) database was performed on the 970 significant DEGs from (D). Enriched GO:BP terms are depicted as bubbles, where color represents significance (*padj*) and bubble size indicates RichRatio (DEGs present in the term divided by the total genes in the term). A plus sign (+) indicates enrichment in the AM + d6 stretch condition, and vice versa for (-). (F) Volcano plot showing significant DEGs between AM + d0/3/6 stretch versus AM alone (log2FC>|0.585|, *padj*<0.05). Key DEGs are named. (G) Pathway analysis using GO:BP was performed on the 927 significant DEGs from (F). Enriched GO:BP terms are depicted as bubbles, where color represents significance (*padj*) and bubble size indicates. A plus sign (+) indicates enrichment in the AM + d0/3/6 stretch condition, and vice versa for (-). (H) Gene set enrichment analysis (GSEA) was performed on a less stringent DEG list (*padj*<0.05) between AM + d0/3/6 stretch and AM alone. All Wnt-related terms in the MSigDB collection of annotated gene sets were interrogated. Normalized enrichment scores (NES) are shown, with bubbles representing the number of DEGs found in a gene set (size) and NES significance (color). (I) Leading-edge genes were calculated from the most significant (FDR<0.25) gene sets (i-iv) and their rank metric scores plotted as a heatmap.

Since these findings support the mechanosensitive and mechanoresponsive nature of intra-articular adipose tissue, we sought to determine whether injurious mechanical loading disrupts adipogenesis, and what mechanisms may regulate this effect. Stromal cells derived from healthy IFP were grown on soft substrate (10 kPa) BioFlex plates, coated with collagen I. Cells were then maintained in growth medium (GM) or in adipogenic medium (AM) to induce differentiation. The AM samples were either kept static or subjected to dynamic mechanical loading using the FlexCell system, in which elastic BioFlex plates permit tensile loading of cell monolayers (Fig 5B). 10% biaxial deformation at 0.5 Hz was undertaken for a period of 4 h on day 6 (terminal differentiation) or throughout differentiation on days 0, 3, and 6. This level of dynamic loading was considered abnormal and injurious, especially for continuous prolonged periods. Cells were harvested for RNA isolation 4 h following the final stretching event, or at the equivalent timepoint for static (unstretched) cells kept in GM or AM. Bulk RNA-seq was then performed, to uncover transcriptional signatures and perturbed functions associated with adipogenic differentiation and with dynamic loading (Fig S5C). Principal component analysis showed clearly separate clustering of GM- and AM-treated cells, and closer but still distinct clustering of AM samples subjected to stretching throughout or only at terminal differentiation (Fig 5C). The first principal component was driven by treatment with GM or AM, and the second component largely by the presence or absence of dynamic loading. 3,787 DEGs (*padj*<0.05, log2FC>|0.585|, FPKM>1) were detected in static AM-compared to GM-treated cells, with notable downregulation of Wnt signaling, and matrix organization and assembly terms (Fig S5D-E, Tables S4-5). Unsurprisingly, adipogenic pathway terms like lipid storage and brown fat cell differentiation were enriched in the AM condition.

Compared to static AM, cells stretched at terminal differentiation had 970 significant DEGs, with genes such as *Acta2* (encoding αSMA), *Piezo2*, *Tgfb1*, *Tnc* and Wnt pathway genes *Lgr6* and *Wnt2* upregulated with stretching (Fig 5D). These DEGs underpinned an enriched hypoxia response and ERK signaling terms in the cells stretched at terminal differentiation (Fig 5E). Cells stretched throughout differentiation on days 0, 3, and 6 had 927 significant DEGs compared to static AM cells, including upregulation of fibrotic markers *Acta2*, *Tnn*, *Tnc* and *Tagln*, and *Wnt2*, and reduced induction of adipogenic markers like *Plin1*, *Adipoq*, *Fabp4* and *Pparg* (Fig 5F). Adipogenic pathway terms were negatively enriched in cells stretched throughout differentiation, while fibroblast proliferation, matrix organization and cell migration were all positively enriched (Fig 5G).

We then performed a two-layered differential expression analysis, separately comparing AM treatment alone or AM with stretching throughout differentiation, to the baseline (GM), in order to capture condition-specific transcriptional responses (Fig S5F-H). In relation to AM treatment alone, stretching at days 0, 3, and 6 broadly suppressed induction of adipogenic genes and elevated expression of fibrotic genes, compared to baseline (Fig S5F-G). AM treatment with stretching also resulted in relative upregulation of genes that promote Wnt signaling, with modest decreases in genes that inhibit Wnt (Fig S5H). The inherent complexity and in-built negative feedback mechanisms of the Wnt pathway then prompted closer analysis of more nuanced shifts in Wnt gene expression programs. For this, we performed gene set enrichment analysis (GSEA) to interrogate all Wnt-related gene sets available on the Molecular Signatures Database (MSigDB) of annotated mouse gene sets (Fig 5H). A less stringent list of DEGs between AM treatment with stretching throughout differentiation and AM alone was used (*padj*<0.05). Terms favoring promotion of Wnt signaling were enriched, whereas terms associated with negative regulation of the Wnt pathway registered negative normalized enrichment scores. The leading-edge genes – genes that contribute most significantly to driving enrichment scores – were extracted from significant positive gene set terms (FDR>0.25; Fig 5I i-iv), and included Wnt ligands, alongside Notch pathway genes. Together, these results demonstrate an inhibitory effect of persistent injurious loading on adipogenesis of IFP-derived stromal cells and implicate upregulated Wnt/β-catenin signaling as a putative mediator in this process.

### Wnt signaling overactivation diminishes intra-articular adipose signature

Given that persistently elevated canonical Wnt/β-catenin signaling is reported in OA joints^38,39,42,72^, combined with our findings above supporting an anti-adipogenic, pro-fibrotic role for Wnt, we sought to determine the role of chronic overactivation of the Wnt/β-catenin pathway on joint adipose tissue *in vitro* and *in vivo*. We first took the IFP from joints of naïve mice and cultured them as explants for 48 h. During this time, they were treated with the potent Wnt/β-catenin activator CHIR99021, combinatorial WNT3A and R-spondin 2, or their respective vehicle controls (Fig 6A-F, Fig S6A-B). CHIR99021, which stabilizes β-catenin by inhibiting glycogen synthase kinase-3, strongly induced Wnt activity in IFP explants, illustrated by upregulation of Wnt target genes *Axin2*, *Lef1*, and *Tcf3* (Fig 6B). Alongside induction of Wnt signaling, CHIR99021 suppressed expression of the lipid droplet-associated gene *Plin1*, and induced the fibrotic genes *Acta2* and *Fbln2* (Fig 6C, Fig S6A). We then sought to activate Wnt signaling using the prototypical Wnt ligand WNT3A, which engages Frizzled/LRP receptor complexes to promote Wnt/β-catenin signaling, and its agonist R-spondin 2, which we have previously shown is secreted after joint injury, promoting pathological crosstalk in joint tissues^37^. Incubating IFP explants with WNT3A and R-spondin 2 induced *Axin2* and *Lef1*, in parallel with suppression of adipogenic genes and corresponding upregulation of fibrotic markers (Fig 6E-F, Fig S6B).

**Fig 6.**
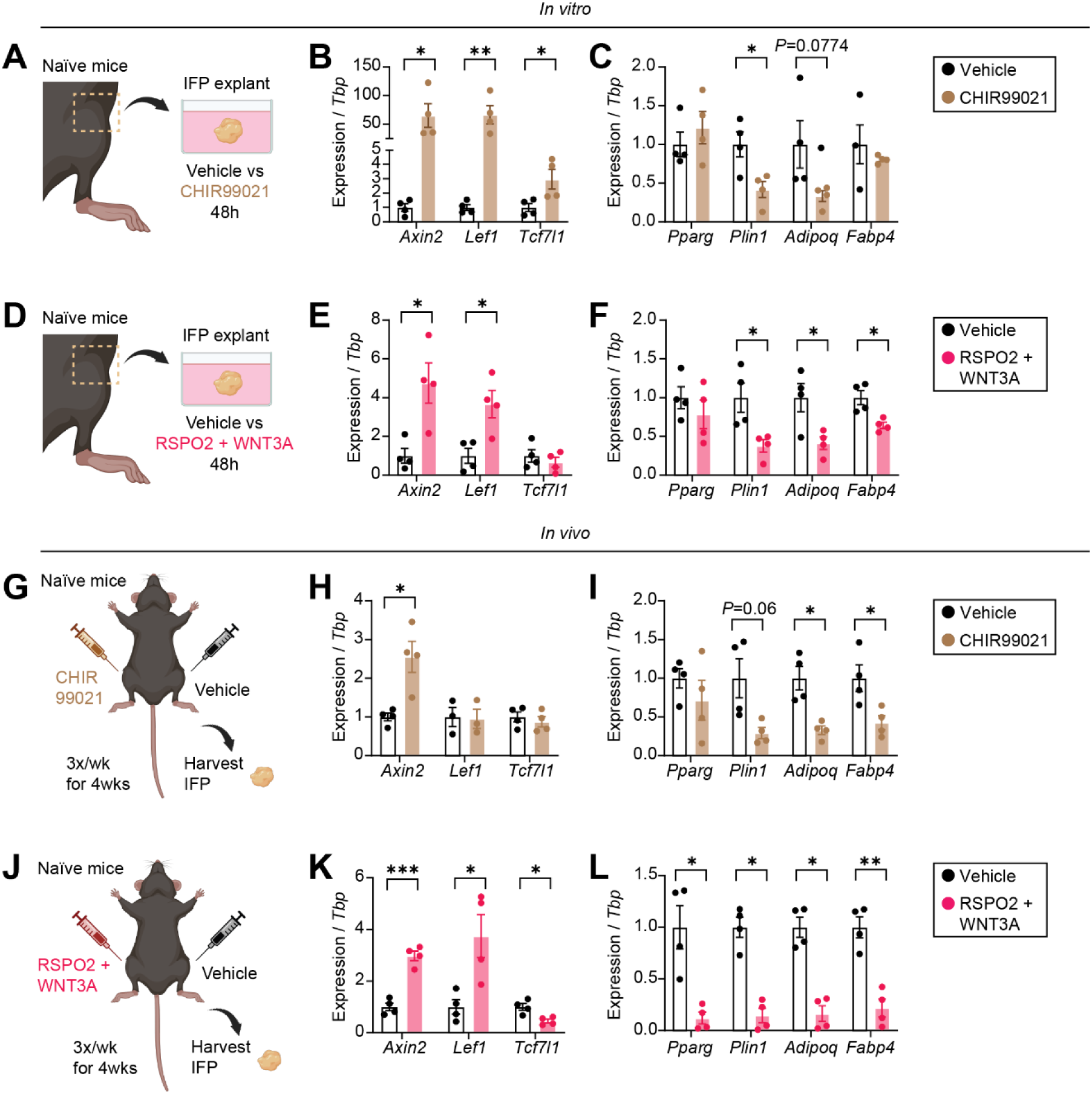
Wnt signaling overactivation diminishes intra-articular adipose signature. (A-F) IFP explants from naïve mice were cultured in adipogenic maintenance medium for 48 h in the presence of 5 µM CHIR99021 or DMSO vehicle (A-C), or 200 ng/mL RSPO2 + 20 ng/mL WNT3A or PBS vehicle (D-F) (n=4). Relative expression of Wnt pathway genes (B, E) and adipogenic genes (C, F) was assessed. (G-L) Naïve mice received thrice-weekly injections for four weeks, directly into the IFP. 5 µg of CHIR99021 or 5% DMSO/95% PBS vehicle (G-I), or 500 ng RSPO2 + 50 ng WNT3A or PBS vehicle (J-L) were delivered in a total volume of 5 µL per injection (n=4). Treatments and their respective vehicles were injected into paired right and left limbs of the same mouse. Relative expression of Wnt pathway genes (H, K) and adipogenic genes (I, L) was assessed. For (B-C and E-F), unpaired two-tailed t-tests were performed, where \**P*<0.05, \*\**P*<0.01. For (H-I and K-L), paired two-tailed t-tests were performed, where \**P*<0.05, \*\**P*<0.01, \*\*\**P*<0.001. For (B-C, E-F, H-I, and K-L), transcript levels were normalized to *Tbp* housekeeper expression, vehicle groups for each gene were set to 1, and error bars are means±SEM.

Then, to extend these findings to an *in vivo* setting, we injected CHIR99021 or R-spondin 2/WNT3A directly into the IFP of naïve mice thrice weekly over the course of 28 days (Fig 6G-L, Fig S6C-E). This sought to test whether overactive Wnt/β-catenin signaling, as observed in OA joint tissues, is sufficient by itself to induce anti-adipogenic effects *in vivo*. Vehicle was administered to the paired contralateral knee. After harvesting IFP tissues, expression of Wnt, adipogenic, and fibrotic genes were assessed. CHIR99021 administration showed elevated *Axin2*, alongside downregulation of adipogenic genes and upregulation of fibrotic markers, compared to vehicle-injected IFP (Fig 6G-I, Fig S6D). IFP repeatedly administered with R-spondin 2/WNT3A showed robust induction of *Axin2* and *Lef1* expression, and very strong suppression of the adipogenic genes *Pparg*, *Plin1*, *Adipoq*, and *Fabp4*, while the fibrotic marker *Col1a1* was upregulated (Fig 6J-L, Fig S6E). Together these data support the notion that overactive Wnt/β-catenin signaling, as seen in OA, diminishes the adipogenic signature of intra-articular adipose tissue and results in concurrent onset of fibrotic hallmarks.

## Discussion

There is a growing focus on the role of adipose tissue in joint diseases like OA, yet little is known about the physiological and pathological functions of local, intra-articular adipose tissue. Here we characterized the spatiotemporal dynamics of fibrotic joint adipose atrophy in PTOA, demonstrating a convergent role for overactive Wnt/β-catenin signaling and mechanical loading in this process.

Having shown that joint injury induced a progressive loss of intra-articular adiposity, we sought to determine the mechanisms underpinning this phenomenon. While stromal progenitors from the synovium and IFP of healthy mice underwent normal adipogenic differentiation *in vitro*, comparable to those derived from WAT, cells from PTOA joints had an impaired capacity for adipogenesis. The Wnt/β-catenin pathway is known to be anti-adipogenic and pro-fibrotic^43,47,69,70^, and overactive canonical Wnt signaling is a feature of OA^38,39,42,72^. Stromal cells cultured from injured joints had higher basal expression of Wnt genes prior to differentiation, possibly predisposing them away from an adipogenic fate, and chronic β-catenin activation during differentiation inhibited adipogenic gene induction and lipid droplet formation. We previously uncovered synovial lining fibroblasts as secretors of the Wnt agonist R-spondin 2 in PTOA^37^, yet the comprehensive identities and sources of Wnt ligands (and agonists) that orchestrate this complex process demand further characterization if we are to effectively target Wnt signaling as a therapy to mitigate fibrosis and its sequelae in OA.

Adipose tissue depots are not typically exposed to dynamic biomechanical perturbation. However, being situated in highly mobile synovial joints places intra-articular adipose tissue under frequent and varied mechanical deformation, representing another unique aspect of this depot. Since the soft tissues in destabilized joints, including synovium and adipose, are susceptible to persistent and abnormal biomechanical forces, we assessed whether injurious dynamic loading would alter adipogenic differentiation of IFP-derived stromal cells. It is important to note that the precise micro-and macro-loading environment of intra-articular adipose tissue is yet to be empirically determined, and what truly constitutes ‘injurious in this setting remains conjecture. Prolonged loading during differentiation had the most inhibitory effect on adipogenic differentiation, instead promoting a matrix-rich, fibroblastic gene program, reminiscent of myofibroblasts rather than adipocytes. Wnt/β-catenin signaling was enriched in this condition, pointing to a convergent mechanism whereby abnormal mechanotransduction after injury promotes overactive Wnt activity, in turn impairing *de novo* adipogenesis and contributing to intra-articular adipose atrophy.

While adipose distribution in joints like the knee has been partly categorized into defined compartments like the IFP, it remains a spatially dispersed depot. This has made studying and quantifying joint adipose tissue a difficult task, and perhaps why studies have predominantly focused on the IFP alone, given its size and somewhat easily defined boundaries. This is particularly relevant in mice, where no other intra-articular adipose tissue in the knee besides the IFP is feasibly accessible for microdissection. Using a 2D histomorphometric approach, we quantified relative intra-articular adipocyte area in the anterior and posterior compartments of the knee based on morphological segmentation of adipocytes. This adipocentric approach showed that relative adipocyte coverage diminished as PTOA progressed, complementing findings from other groups that evaluated joint adipose atrophy from a fibrotic standpoint, using picrosirius red staining^49,50^. Spatial transcriptomics allowed for advanced, high-resolution analysis of adipocytes and fibroblasts within their anatomic context, extending the finding that in PTOA, joint adipose tissue comprised of adipocytes shifts to a more fibrotic profile comprised of activated fibroblasts – linking adipocyte loss to fibrosis onset. This is relevant given that IFP fibrosis correlates to clinical stiffness^75^, which itself is associated with pain and disease severity in OA^23,24,27,28^.

To study intra-articular adipose atrophy in a more quantitative, three-dimensional fashion, we developed a method to volumetrically quantify adiposity throughout the whole knee joint by contrast-enhanced µCT using the lipophilic compound osmium tetroxide. This approach has previously been used for imaging bone marrow adipose tissue^65^, and here it allowed us to visualize the distribution and extent of lipid-laden adipocytes throughout healthy and PTOA mouse joints, corroborating our finding of adipose atrophy in PTOA. This method does not resolve individual adipocytes, and osmium tetroxide does not discriminate between intracellular versus extracellular lipids, but it does provide high resolution volumetric mapping of the intra-articular adipose depot, and analysis of attenuation signal patterns may permit inference about tissue fibrosis onset. In the longer term, a non-destructive approach that allows longitudinal imaging of joint adipose tissue volume and composition would greatly advance our ability to understand the dynamics of adipose loss and fibrosis.

Since clinical studies indicate that intra-articular adipose tissue size is not correlated with BMI^8,18–21^ and that it may have unique fundamental functions, we compared the transcriptional signature of healthy IFP to classical WAT in mice. Pathway and cell type enrichment analyses indicated that the murine IFP was more matrix rich and stromal cell-dominated, plus more highly associated with innervation and neuronal cell types, compared to WAT, which harbored a more enriched hematopoietic niche. This aligns with our understanding of the IFP as a highly neurovascular tissue, whose fibrotic remodeling is clinically associated with pain. There is conflicting evidence^19,20^ regarding whether joint adipose tissue undergoes inflammatory changes seen in classical WAT depots, such as adipocyte hypertrophy, immune infiltration, and macrophage crown-like structure formation. Recent comparative analysis of human intra-articular adipose tissue and subcutaneous adipose tissue from OA patients points to a pathological phenotype of joint adipose tissue in disease, including secretion of pro-inflammatory cytokines that perturbed other joint-resident cell types^76^. With new mouse models emerging that can selectively target the joint adipose depot^10^, there is strong impetus to better characterize the homeostatic and disease-associated functions of intra-articular adipose tissue, and understand its commonalities and distinctions with other peripheral adipose depots.

This study deals primarily with adipogenesis and fibrosis in OA, but inflammation and pain are two additional disease-relevant phenomena whose manifestations are closely intertwined. In addition to the fibrotic phenotype investigated here, work from our group has shown a marked inflammatory and neurotrophic/pro-algesic program inherent to OA synovium and intra-articular adipose tissue in both mice and humans^31,77^. Ongoing and future investigations must acknowledge the holistic, multi-faceted nature of joint disease, and strive to incorporate all these phenomena. Given its highly neurovascular disposition and its potential as a regenerative niche, there is strong impetus to better understand intra-articular adipose tissue ‘quality’ and ‘quantity’, and how it relates to arthritis pathogenesis. We have identified convergent mechanical loading and overactive Wnt/β-catenin signaling as putative mechanisms underpinning atrophy of intra-articular adipose tissue, shedding critical light on its pathophysiological functions in OA, and uncovering a translational target for achieving the ultimate goal of restoring healthy joint function.

## Methods

### Mice

Male and female C57BL/6J mice were procured from Jackson Laboratories (Strain #000664), and all animal procedures were performed in accordance with the University of Michigan’s approved institutional animal care and use protocols. Mice were housed communally on a 12 h light:dark cycle and given *ad libitum* access to food and water. All procedures and harvests took place between 12 and 16 weeks of age.

### Anterior cruciate ligament rupture model of post-traumatic osteoarthritis

Non-surgical joint injuries were performed on 12-week-old mice to induce post-traumatic osteoarthritis (PTOA), as previously published^30,33,35^. Briefly, mice were subjected to inhaled isoflurane anesthesia and received unilateral hindlimb pre-loading and cyclic pre-conditioning, followed by a single mechanical overload resulting in complete anterior cruciate ligament rupture (ACLR). Mice were then given 5 mg/kg carprofen subcutaneously for analgesia. Sham mice were subjected only to anesthesia and analgesia. Non-surgical ACLR procedures were performed using either a modified CellScale Univert S2 or an Electroforce 3300AT mechanical testing system, which have been shown to have high concordance^35^.

### 2D histomorphometry

The anterior and posterior synovium/adipose tissue histomorphometry datasets consist of 5 µm sagittal sections from the medial region of male and female C57BL6/J mouse knee joints, stained with Safranin O/Fast Green and imaged at 20x on a Nikon Eclipse Ni E800 with DS-Ri2 camera. We employed existing sections from our prior study characterizing sex-dependent phenotypes following joint injury using the same ACLR model described above^31^. Samples are grouped based on sex (male or female) and condition (Sham, 7d ACLR, 28d ACLR). Images were excluded from this study if they had substantial tissue shredding or imaging artifact, any portion of the region of interest was cut off in the image, or a large spot from staining covering regions of the intra-articular adipose tissue.

Manual polygonal outlining of the anterior and posterior synovium and intra-articular adipose tissue were performed. To identify adipocytes within masks, a custom MATLAB script was used to manually mark up to ten distinct adipocytes within each image. Images were converted to *Lab* colorspace, and the average pixel lightness (*L*) for adipocytes was calculated, distinguishing it from other tissues and noise. For each image, an initial lightness threshold for adipocytes was defined as the average adipocyte lightness minus 5, and any pixel with a lightness higher than this threshold was classified as an adipocyte. To account for differences in contrast between images, this threshold was dynamically relaxed by the user, in blinded fashion, until adipose tissue was comprehensively captured. Any remaining defects in the tissue outline following thresholding were user-corrected by manual polygonal outlining, if present.

Total outline area (synovium and adipose tissue) and adipose only area (adipocytes) were calculated. The adipocyte fraction (percent area identified as adipose tissue containing adipocytes) was calculated by dividing the adipocyte-only area by the total area for each image. Technical replicates from different sections of the same mouse were averaged to yield adipocyte fractions for each individual mouse (biological replicates) then one-way ANOVA with post-hoc Tukey testing was used to compare values between Sham, 7d ACLR and 28d ACLR.

### Osmium tetroxide staining and imaging

#### Osmium tetroxide processing

Male C57BL6/J mice were subjected to unilateral ACLR at 12 weeks of age. After 28 days, whole left (contralateral) and right (28d ACLR) hindlimbs were harvested and fixed in 10% neutral-buffered formalin for 24 h. Limbs were then directly transferred to osmium tetroxide staining solution (1% final concentration) that was diluted in Sorensen’s Phosphate Buffer for a total volume of 20 mL per sample. Hindlimbs were stained for 48 h then washed with Sorensen’s Phosphate Buffer before μCT scanning.

#### Micro-computed tomography (μCT) scanning and analysis

μCT scans were acquired using a Bruker 1176 μCT imaging system. Final scans were acquired at a 9 μm isotropic voxel resolution, 0.25-degree rotation step, 4 frame averages, and 360-degree scan. Segmentation and quantification were performed in Dragonfly 3D World (version 2024.1 Build 1627). Scans were loaded into the software and manual outlines were drawn on 2D axial slices to equally segregate the anterior and posterior region of the articular space. Superior and inferior boundaries were established at the physis of each long bone using the 2D sagittal view. A 20,000+ arbitrary unit (AU) threshold on a 16-bit scale was empirically determined and set to allow for accurate segmentation of contrast-enhanced features bound by osmium tetroxide while minimizing false detection of non-ROI structures (e.g. bone, soft tissue, background). Manual outlines and fat signal thresholds were then intersected, and volume and attenuation data were exported. Representative images were generated from a cleaned 3D rendering, with extra-articular tissue removed to visualize only intra-articular structures. Attenuation values were converted from AU to Hounsfield Units (HU) using calibration data from a scan of a phantom consisting of two hydroxyapatite concentrations^31^.

#### Post-scan decalcification and histology

Following μCT image acquisition and quality control, limbs were decalcified in 100% Immunocal (formic acid) for 12 h. After decalcification, limbs were subjected to standard paraffin processing then embedded in paraffin blocks that were sagittally sectioned at 5 μm thickness. Slides were mounted with CytoSeal and coverslipped, then brightfield-imaged using a Keyence BZ-X810 microscope with a 10x objective, to assess osmium tetroxide penetration into intra-articular adipose tissue and to visualize staining throughout intra-articular tissues.

### Intra-articular IFP injections in mice

Our standard intra-articular injections are 2 mm in depth, and deliver injectate through the patellar tendon and IFP, directly into the intra-articular cavity (synovial fluid). Based on calculations of the depth of the IFP in the knee, using historical μCT scans of mouse knee joints, we determined that a 1.25 mm depth injection would directly deliver injectate into the IFP. Comparison of macroscopic trypan blue staining following injection at both depths showed 1.25 mm to be suitable for direct IFP injections (Fig S6C).

### Overactivation of Wnt signaling in the IFP

Naïve 12-week-old C57BL6/J male and female mice were given thrice-weekly injections into the IFP, for four weeks. 33-gauge needles were used to deliver 5 µL per injection, with an injection depth of 1.25 mm. For the first experiment, left knees received vehicle (5% DMSO/95% PBS) and right knees received the β-catenin stabilizer CHIR99021 (5 µg per injection, in 5% DMSO/95% PBS), to chronically activate canonical Wnt signaling. For the second experiment, left knees received vehicle (PBS) and right knees received the Wnt ligand WNT3A (50 ng per injection, in PBS) and the Wnt signaling agonist R-spondin 2 (RSPO2, 500 ng per injection, in PBS), to chronically activate canonical Wnt signaling. After four weeks, the left and right IFPs were dissected and immediately homogenized in TRIzol using a Bullet Blender on speed 8 for three pulses of 30 s, before proceeding to RNA isolation.

### Cell culture and adipogenic differentiation

As described previously^78^, WAT was harvested, minced, then digested in Collagenase D (1.5 U/mL), Dispase II (2.4 U/mL), and 10 mM CaCl_2_ for 15-20 min in a 37°C water bath with agitation. Digested tissues were filtered through a 100 μm strainer and centrifuged at 500 x *g* for 5 min to pellet cells (the stromal-vascular fraction, SVF). The SVF cell pellet was resuspended and passed through a 40 μm strainer and centrifuged again. The pellet was resuspended in DMEM/F12 GlutaMAX containing 10% FBS and antimycotic/antibiotic then seeded onto tissue culture-treated plates. Cells were expanded in culture up to three passages. For adipogenic differentiation, cells at confluency were exposed to adipogenic induction medium comprised of DMEM/F-12 GlutaMAX with 10% FBS, antimycotic/antibiotic, 0.5 μg/mL insulin, 5 μM dexamethasone, 1 μM rosiglitazone and 0.5 mM IBMX. Three days after induction, cells were switched to a maintenance medium containing 10% FBS, antimycotic/antibiotic, and 0.5 μg/mL insulin, which was replaced every two days.

Stromal cells from mouse synovium/IFP, often termed fibroblast-like synoviocytes or cultured synovial fibroblasts, were isolated and cultured as described previously^37,79^. Briefly, knee synovium and IFP from healthy mice or mice subjected to 7d ACLR was harvested and digested in DMEM containing 400 µg/mL DNaseI, 400 µg/mL Liberase TM, and 400 µg/mL Collagenase IV for 30 min, followed by centrifugation at 500 x *g* for 5 min. The pellet was resuspended in DMEM containing 10% FBS and antimycotic/antibiotic, then cells were seeded onto CellBind plates to promote adhesion. Cells were expanded in culture up to three passages. Adipogenic induction and maintenance was performed as above for WAT SVF. For initial evaluation of adipogenic differentiation of IFP-derived stromal cells, adipogenic medium (induction and maintenance medium, as above) was compared to an alternative media combination, 3T3L1, comprised of induction medium (DMEM/F-12 GlutaMAX with 10% FBS, antimycotic/antibiotic, 5 μg/mL insulin, 1 μM dexamethasone, and 0.5 mM IBMX) followed by maintenance medium (DMEM/F-12 GlutaMAX with 10% FBS, antimycotic/antibiotic, 5 μg/mL insulin).

To modulate Wnt signaling, cells were treated with vehicle (PBS), 200 ng/mL recombinant mouse R-spondin 2 (RSPO2, in PBS), 20 ng/mL recombinant mouse WNT3A (in PBS), or a combination of the two, either for the duration of adipogenic differentiation (fresh treatments with every media change) or for the final 24 h of differentiation. Alternatively, cells were treated with vehicle (DMSO) or 5 μM CHIR99021 (in DMSO) throughout differentiation.

### Mechanical loading of cells

IFP-derived stromal cells were seeded onto CellSoft BioFlex plates (10 kPa stiffness) coated with collagen I (FlexCell International). At confluency, cells were either subjected to adipogenic differentiation as described above or kept in growth medium. For mechanical loading, cells were placed in the FX-6000T Tension System, within a 37°C CO2 incubator. Cells were subjected to a regimen of 10% biaxial strain at 0.5 Hz for a duration of 4 h, either on days 0, 3, and 6 of differentiation, or just on day 6. All cells were harvested in TRIzol on day 6, 4 h after the cessation of loading.

### IFP explant treatments

The IFP of both knee joints was dissected from naïve 12-week-old male and female C57BL6/J mice, then placed directly into wells of a 96-well plate containing adipogenic maintenance media (DMEM/F-12 GlutaMAX with 10% FBS, antimycotic/antibiotic, 5 μg/mL insulin). Explants were treated for 48 h in the presence of 5 μM CHIR99021 or vehicle (DMSO), or RSPO2 (200 ng/mL) + WNT3A (20 ng/mL) or vehicle (PBS). Following treatment, IFP explants were homogenized in TRIzol using a Bullet Blender on setting 8, with 30 s pulses, prior to RNA isolation by phenol chloroform extraction, and gene expression analysis as described below.

### Oil Red O staining

Fresh Oil Red O working solution was made using three parts 0.5% Oil Red O in isopropanol, to two parts PBS, then particulates were filtered out. Media on cells was aspirated then 4% paraformaldehyde fixative was added for 1 h at room temperature. Fixative was then removed, cells were washed with PBS, and Oil Red O working solution was added for 1 h at room temperature. After this, Oil Red O stain was removed, and cells were washed two to three times with PBS, prior to imaging. A Lionheart FX (BioTeK) was used for phase contrast imaging of cells prior to Oil Red O staining, and for fluorescence imaging of Oil Red O stain using the Texas Red channel.

### Gene expression analysis

Cells were lysed and homogenized in TRIzol then RNA was isolated by phenol chloroform extraction. IFP was homogenized in TRIzol using a BulletBlender on setting 8 for 30 s pulses until the tissue was fully homogenized, then RNA isolated by phenol chloroform extraction. cDNA was synthesized using the random primers method with a High-Capacity cDNA Reverse Transcription Kit. PowerSYBR chemistry was used for quantitative real-time PCR (qPCR). For analysis, expression was normalized to levels of an internal housekeeping gene, *Tbp*, using the 2^-Δ^ ^ΔCt^ method. Primer sequences can be found in Table S3.

### Protein isolation and Western blotting

Stromal cells from WAT or IFP were cultured in growth medium or adipogenic medium. Nuclear protein extractions were performed as reported previously^80,81^. Whole cell protein lysate was isolated using radioimmunoprecipitation buffer as done previously^82^, from mouse bone marrow-derived macrophages grown in DMEM containing 10% FBS, antibiotic/antimycotic, L-glutamine, and 10 ng/mL M-CSF. Protein extracts were quantified via colorimetric bicinchoninic acid assays then 15 µg of protein was mixed with Bolt LDS Sample Buffer (4X) and Bolt Sample Reducing Agent (10X), then heated at 70°C for 10 min. Protein was loaded onto a 12-well 4-12% bis-tris polyacrylamide gel, alongside a PageRuler Prestained Plus rainbow marker as a size reference. Proteins were transferred onto nitrocellulose membranes using the iBlot 3 Dry Blotting System with the preprogrammed “Low Molecular Range” transfer method. The membrane was blocked in 5% bovine serum albumin in TBST for 1 h and incubated with primary antibodies against PPARɣ (CST, #2443S; 1:1,000) and β-actin (Sigma, A5316; 1:5,000) overnight at 4°C followed by incubation with HRP-conjugated goat anti-rabbit (Thermo #31460; 1:10,000) or goat anti-mouse (Thermo #31430; 1:10,000) secondary antibodies for 1 h at room temperature. Signal was developed using SuperSignal™ West Pico PLUS substrate and imaged on an Invitrogen iBright CL1500 Imaging System.

### Bulk RNA-sequencing

For transcriptional comparison of WAT and intra-articular adipose tissue, naïve male and female C57BL/6 mice were euthanized at 12 weeks of age. The inguinal subcutaneous WAT was harvested and the inguinal lymph node removed; then the IFP from both knees was harvested. The IFP was used since it is the only feasibly accessible adipose depot in the murine knee joint, for dissection.

Tissues were immediately placed in RLT buffer in Precellys CK28 soft tissue homogenization tubes and kept on ice. Homogenization was performed using a Precellys Evolution at 4°C using the built-in soft tissue setting once (IFP) or thrice (IWAT). Qiagen RNeasy kits (Micro for IFP, Mini for IWAT) were used for RNA isolation, including DNase treatment to remove carryover genomic DNA. RNA was submitted to the University of Michigan Advanced Genomics Core for quantification and quality control on an Agilent Bioanalyzer, then cDNA libraries were prepared using low input kits (Takara SMART-Seq v4 PLUS) and subjected to 151 bp paired-end sequencing on an Illumina NovaSeq. BCL Convert Conversion Software (v4.0) was used to generate de-multiplexed Fastq files.

Snakemake^83^ was used to manage the bioinformatics workflow in a reproducible manner. Reads were trimmed using Cutadapt (v2.3) and evaluated with FastQC (v0.11.8) to determine data quality. Reads were then mapped to the reference genome GRCm38 using STAR (v2.7.8a) and assigned count estimates to genes with RSEM (v1.3.3). Quality control metrics from several different steps in the pipeline were aggregated by multiQC (v1.7). The biomaRt R package was used to import chromosome name and gene biotype information for each gene. For sequencing data filtering and processing, count matrices were filtered to keep genes with at least 10 reads and genes with an FPKM of at least 1, in at least n samples (n = min number of samples in any group). Non-protein-coding genes and sex-linked genes were also filtered out.

For transcriptomic profiling of IFP-derived stromal cells subjected to adipogenic differentiation and mechanical loading, cells were isolated, cultured, differentiated, and treated as per above. TRIzol was added to sample wells 4 h after the completion of the final stretching event, to harvest and lyse cells. Phenol chloroform extraction followed by RNA isolation using a Qiagen RNeasy Mini Kit with DNase digestion was performed to isolate RNA, before submission to Novogene for bulk RNA-seq. Paired-end sequencing of polyA-enriched mRNA libraries was done on an Illumina NovaSeq X Plus, and raw Fastq files were aligned to the GRCm38 reference genome using HISAT2^84^. FeatureCounts (v2.0.6) was used to count the read numbers mapped to each gene. Count matrices were filtered to keep genes with at least 10 reads and genes with an FPKM of at least 1, in at least n samples (n = min number of samples in any group). Non-protein-coding genes and sex-linked genes were filtered out.

### Transcriptomic analyses

Analysis of bulk RNA-seq data was undertaken in R. Principal component analysis (PCA) using pcaExplorer^85^ assessed sample clustering and outlier status, based on 95% confidence interval. Analysis of differentially expressed genes (DEGs) was performed using DESeq2^86^, and significant DEGs were defined as adjusted p-value (*padj*)<0.05 and log2FC>|0.585|, unless otherwise stated. Volcano plots of DEGs were generated in GraphPad Prism 10, and DEG heatmaps were generated in Prism or in R using pheatmap. To analyze enriched pathways and biological functions, significant DEG lists were input into PantherDB^87^ for statistical enrichment testing using the Gene Ontology (GO) Biological Processes (BP) or Molecular Function (MF) aspects, or Reactome. Outputs were corrected for False Discovery Rate (FDR) and terms were manually categorized into broad biological themes.

Bubble plots for pathway analyses were generated using ggplot2^88^ in R, where color indicates significance (*padj*) and size indicates RichRatio, which is the number of DEGs present in the term divided by the total genes in the term. Cell type enrichment analysis of IFP and WAT was performed using WebGestalt 2024^89^. Significant DEGs (*padj*<0.05 and log2FC>|0.585|) in the positive and negative direction were used as separate inputs, then over-representation analysis was run to identify enriched cell types in each tissue using the Mouse Cell Atlas functional database for consultation, the *Mus musculus* genome as the reference gene set, and with Benjamini-Hochberg FDR correction to calculate *padj*. The top 10 hits were collected and enrichment ratios were calculated as follows: enrichment ratio = (number of genes in the functional database gene set * number of genes in input list) / total number of genes in the reference genome list. For gene set enrichment analysis (GSEA), GSEA desktop software (v4.4.0) was used^90^. A ranked list of DEGs (*padj*<0.05) from the adipogenic medium + days 0/3/6 stretching versus adipogenic medium alone comparison was used as input to run GSEA with default settings, on all annotated gene sets from MSigDB containing the term ‘Wnt’.

Normalized enrichment scores were calculated then visualized as bubble plots using ggplot2. Leading edge genes driving significantly enriched gene sets (FDR<0.25) were determined, and their rank metric scores plotted as a heatmap. Two-layered differential expression analysis was performed to compare curated gene sets (adipogenesis, fibrosis, Wnt signaling) between growth medium and treatment with adipogenic medium alone, or adipogenic medium plus stretching at days 0/3/6. For each gene set, log2FC values from the relevant comparisons were extracted and assembled into a gene-by-condition matrix. Heatmaps were generated using a log2FC color scale, with gray indicating comparisons that were not statistically significant.

For analysis of published bulk RNA-seq datasets, R was used. Data from Sham and ACLR mouse synovium was derived from GSE271903^31^, and heatmaps were generated in GraphPad Prism 10 using log2FC data from DESeq2 analysis. For analysis of published single-cell RNA-seq (scRNA-seq) datasets, the Seurat v5^91^ package in R was used. Data from OA human synovium/IFP was derived from GSE216651^58^ and pseudo-bulk DEG analysis performed using the FindMarkers function. Data from high and low pain human OA synovium was derived from GSE248453^92^ and pseudo-bulk DEG analysis was performed using the FindMarkers function.

### Xenium spatial transcriptomics

Male and female C57BL/6J mice were subjected to Sham or ACLR at 12 weeks of age then harvested one week later for spatial transcriptomics of knee joints. Tissue harvests and processing were performed according to our published protocol^93^. Under inhaled isoflurane anesthesia, mice were transcardially perfused with 20 mL of cold PBS to remove excess blood, followed by 50 mL of cold Z-fix at a flow rate of 200 mL/h. Hindlimbs were disarticulated at the hip joint and fixed for an additional 48 h in 10 mL of Z-fix while nutating at 4ᵒC. After fixation, samples were decalcified with 80% Immunocal (diluted in PBS) for 16 h while nutating at 4ᵒC, followed by paraffin processing in the Michigan Integrative Musculoskeletal Health Core Center’s Structure, Composition, and Histology Core. Samples were embedded with medial side down, and excess wax surrounding the tissues was removed to optimize RNA isolation. Five 20 µm sections per sample were collected for RNA isolation using the Qiagen RNeasy FFPE kit. Due to the large sample size, double the reagents recommended in the protocol were used to effectively remove residual paraffin that might interfere with RNA purity. Isolated RNA was submitted to the University of Michigan Advanced Genomics Core for RNA quality control using the Agilent Bioanalyzer to evaluate concentration, RIN, and DV200.

For mounting onto a 10x Genomics Xenium slide, proximal femur, distal tibia, along with posterior muscles were first removed from the samples. Six hindlimbs were then co-embedded into a single tissue mold (10 x 24 x 5 mm); one multi-tissue section was collected onto a Xenium slide and submitted to the University of Michigan Advanced Genomics Core for spatial transcriptomics using the 10x Genomics Xenium platform. The Mouse Multi-Tissue Atlassing panel was used to detect 379 pre-designed genes, with added multimodal cell segmentation containing antibodies identifying the cell membrane, cytoplasmic RNA, cytoplasmic protein, and nuclei.

The percentage of high-quality gene transcripts (Q-value ≥ 20) were obtained from the Xenium Analyzer pipeline. Xenium Explorer (v3.2.0) was used for sample alignment, cell and transcript overlay, and image export. Each Xenium assay was accompanied by post-processing Hematoxylin and Eosin (H&E) staining. H&E images corresponding to each sample were imported into Xenium Explorer for alignment with Xenium morphology image through at least 100 key points. The intra-articular space and anterior synovium were manually outlined using the aligned H&E in Xenium Explorer. Coordinates from cells within this region were exported and used to computationally subset cells in R (v4.5.0). Cells within the intra-articular compartment from Sham and 7d ACLR joints were integrated via SCTransform normalization and integration (Seurat v5). Non-linear dimensionality reduction using uniform manifold approximation and projection (UMAP) was performed with dims = 1:20 and res = 0.28. Gene marker analysis was performed to identify cell types. DEGs between clusters were identified using the FindAllMarkers function, with *padj*<0.05 and log2FC>|0.585|. Representative markers of the clusters were visualized using violin plots. Cell clusters were exported from R with their respective cell IDs and imported into Xenium Explorer.

## Acknowledgements Funding

This work was directly supported by the US National Institutes of Health (NIH)/National Institute of Arthritis and Musculoskeletal and Skin Diseases (NIAMS) (R01AR080035 to TM) and the Ralph and Marian Falk Medical Research Trust (Catalyst Award to TM). AJK was further supported by NIH/NIAMS (K99AR081894 and R00AR081894), a Pioneer Fellowship from the University of Michigan Medical School, a RAC Grant from the University of Michigan Department of Orthopaedic Surgery, and an Arthritis National Research Foundation Grant (1469667). KDH was further supported by R01DE030716. ELS was further supported by R01DK132073. TM was further supported by NIH/NIAMS (R21AR080502, R21AR082016, R21AR076487, UC2AR082186) and the Department of Defense Congressionally Directed Medical Research Programs (CDMRP) (HT94252310327). We thank the University of Michigan Advanced Genomics Core and Michigan Integrative Musculoskeletal Health Center (P30AR069620).

## Author contributions

AJK conceptualized the project, supervised the project’s execution, designed and performed experiments, analyzed data, and wrote the manuscript. TM conceptualized the project, supervised the project’s execution, and wrote the manuscript. KDH and ELS supervised the project’s execution. All other authors designed and performed experiments, and analyzed data.

## Competing interests

TM is a consultant for Relation Rx. No other authors have competing interests to declare.

## Data, code, and materials availability

All published bulk and single-cell RNA-sequencing datasets are publicly available from NCBI GEO (GSE271903, GSE216651, GSE248453). Requests for newly generated bulk RNA-sequencing and spatial transcriptomic datasets will be considered upon reasonable request to the corresponding authors once this work has undergone peer reviewed publication.

## List of Supplementary Materials

Figs. S1 to S6. Tables S1 to S5.

