## Supplementary Materials for "Wnt/β-catenin signaling regulates fibrotic atrophy of intra-articular adipose tissue in post-traumatic osteoarthritis"

Knights et al, 2026

**Supplementary Figures**

Fig S1. Assessment of joint adipose loss by 2D histomorphometry. Related to Fig 1.

Fig S2. Assessment of joint adipose tissue loss by spatial transcriptomics. Related to Fig 1.

Fig S3. Bulk RNA-seq analysis of IFP and white adipose tissue. Related to Fig 3.

Fig S4. Wnt signaling modulation during adipogenesis. Related to Fig 4.

Fig S5. Mechanical loading blocks adipogenesis. Related to Fig 5.

Fig S6. Chronic Wnt overactivation diminishes intra-articular adipose signature. Related to Fig 6.

**Supplementary Tables**

Table S1. Fat pad vs WAT DEGs

Table S2. Fat pad vs WAT pathway terms

Table S3. Primers

Table S4. DEGs from dynamic loading experiments

Table S5. GO terms from dynamic loading experiments

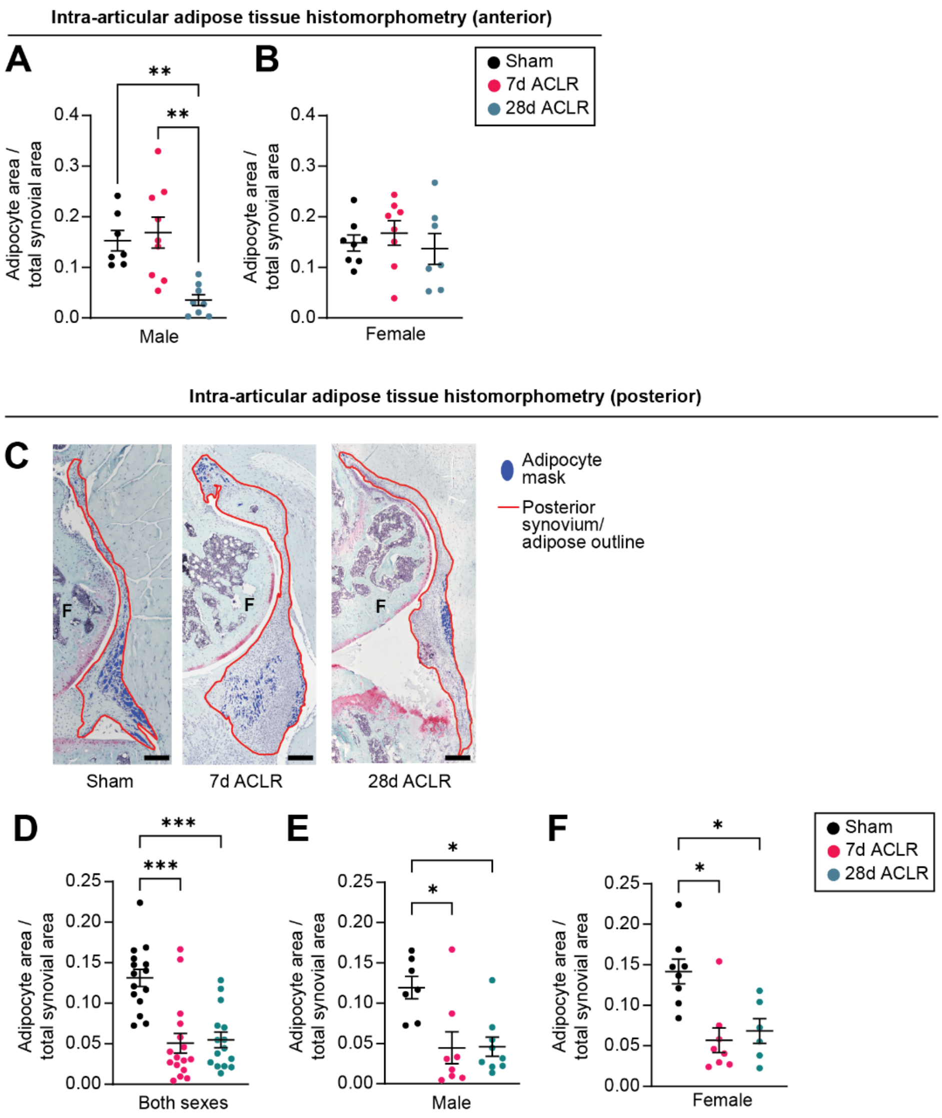


**Fig S1. Assessment of joint adipose loss by 2D histomorphometry. Related to Fig 1.** (A-B) Ratio of anterior adipocyte area to total synovial outline area in Sham, 7d ACLR or 28d ACLR for (A) male (n=7-9 mice) and (B) female (n=7-8 mice), separately. (C-F) Representative images of posterior synovium/adipose tissue from Sham, 7d ACLR, and 28d ACLR sagittal knee joint sections stained with Safranin O/Fast Green (C). Synovium/adipose tissue is outlined in red; adipocyte mask is blue. Ratio of posterior adipocyte area to total synovial outline area in Sham, 7d ACLR or 28d ACLR for (D) both sexes (n=15-16 mice), (E) males only (n=7-9 mice), or (F) females only (n=6-8 mice). One-way ANOVA testing with Tukey's multiple comparisons test (A-B) or Kruskal-Wallis testing with Dunn's multiple comparisons test (D-F) were performed where *$P$<0.05, **$P$<0.01, ***$P$<0.001. Errors bars are means±SEM for (A-B, D-F). ACLR: anterior cruciate ligament rupture. F: femur. Scale bars: 100 μm.

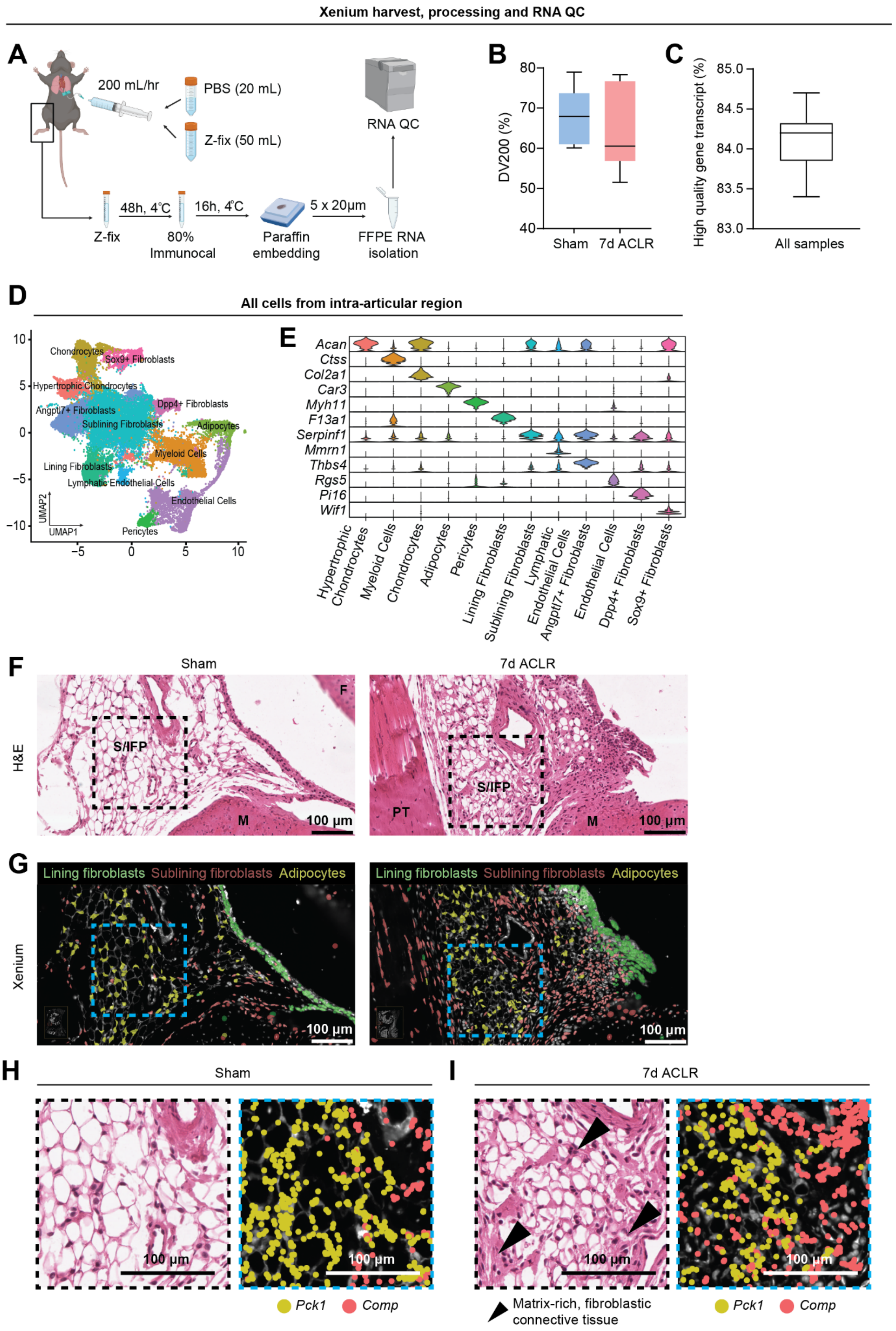


**Fig S2. Assessment of joint adipose tissue loss by spatial transcriptomics. Related to Fig 1.** (A) Schematic showing harvest and processing of Sham and 7d ACLR hindlimbs for Xenium spatial transcriptomics. (B) DV200 (%) of RNA harvested from joint tissue sections processed for spatial transcriptomics (n=6 mice). (C) Percentage of high-quality transcripts (Q-score≥20) from sequencing of all Xenium samples (n=6 mice). (D) Semi-supervised UMAP clustering of intra-articular cell types derived from spatial transcriptomics. (E) Gene markers supporting the identity of each cluster. (F-G) H&E staining of sagittal knee joint sections from Sham (left) or 7d ACLR (right) female mice (F), focused on the anterior synovium and IFP. (G) Xenium spatial transcriptomics showing clustering of synovial lining fibroblasts, sublining fibroblasts, and adipocytes in the synovium and intra-articular adipose tissue from the same sections as in (F). (H-I) Magnified fields of view from (F-G), demarcated by black (H&E) or blue (Xenium) dashed boxes. *Pck1* and *Comp* transcript expression is overlaid for both Sham (H) and 7d ACLR (I) Xenium views, with black arrowheads indicating emergent fibroblastic, matrix-rich tissue regions. For box and whisker plots (B-C), the min, max, median, and interquartile range are shown. PT: patellar tendon. S/IFP: synovium/IFP. M: meniscus. F: femur. Scale bars: 100 μm.

**A**

| Filtering step | Number of genes |
|---|---|
| Pre-filtering | 55,492 |
| Remove duplicate and zero-read genes | 55,372 |
| Min FPKM >1 and Min Count >10 filters | 14,984 |
| Remove non-protein coding genes | 13,027 |
| Remove sex-linked genes | 13,023 |

**B**

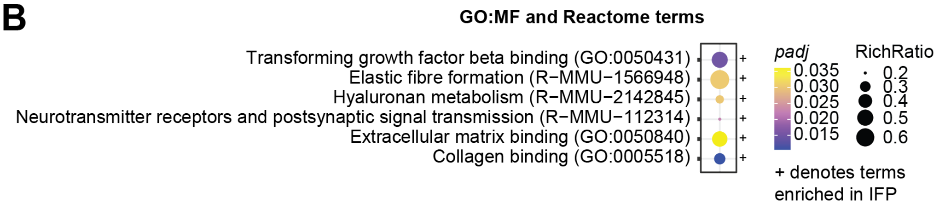


**Fig S3. Bulk RNA-seq analysis of IFP and white adipose tissue. Related to Fig 3.** (A) Gene filtering steps prior to RNA-seq analysis. (B) Enriched parent terms from Gene Ontology: Molecular Function (GO:MF) and Reactome pathway analysis and their tissue-type directionality (+ or -). Bubble color represents significance (*padj*) and bubble size indicates RichRatio (DEGs present in the term divided by the total genes in the term).

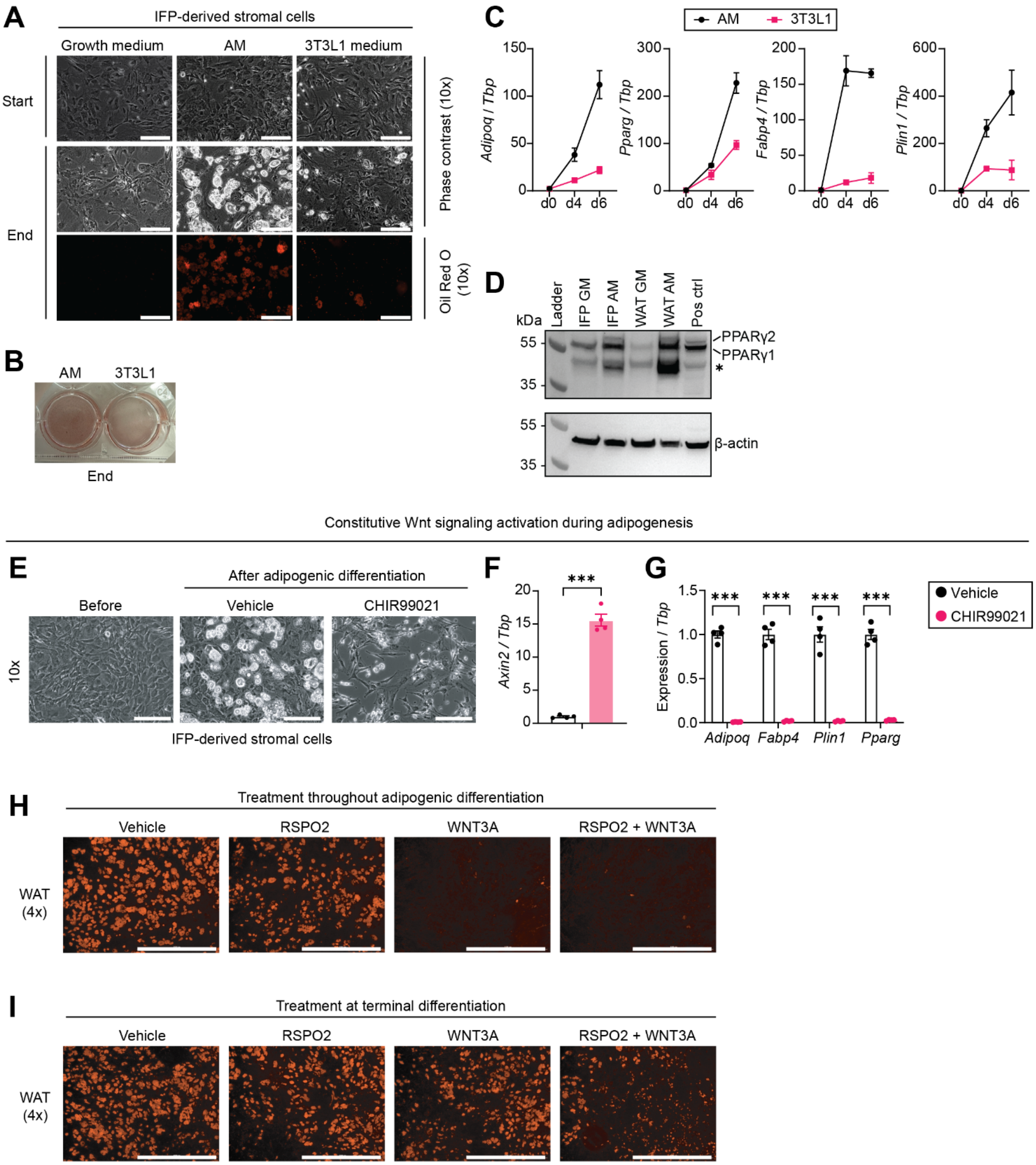


**Fig S4. Wnt signaling modulation during adipogenesis. Related to Fig 4.** (A-C) Stromal cells from healthy mouse IFP were subjected to growth medium or two alternative types of adipogenic differentiation media, adipogenic medium (AM) or 3T3L1 medium. (A) Representative 10x phase contrast images before (top) or after (middle) differentiation, with Oil Red O lipid droplet staining (10x) after differentiation (bottom), for each condition. Scale bars: 200 µm. (B) Macroscopic visualization of Oil Red O staining for the AM and 3T3L1 conditions after differentiation. (C) Expression of adipogenic genes at days 0, 4, and 6 of Adipo or 3T3L1 medium (n=3-4). (D) PPARɣ protein levels in the nuclear extract of stromal cells from WAT or IFP cultured in growth medium (GM) or adipogenic medium (AM). Protein size markers are indicated and PPARɣ isoforms 1 and 2 are shown. The positive control lane is protein extract from bone marrow-derived macrophages. Asterisk indicates background bands/degradation products. (E-G) Stromal cells from healthy IFP were imaged by 10x phase contrast (E) before (left) or after (middle, right) adipogenic differentiation, in the presence or absence of the Wnt signaling activator CHIR99021 (5 µM). Scale bars: 200 µm. After differentiation, expression of *Axin2*

(F) and adipogenic genes (G) was assessed (n=4). Unpaired two-tailed t-tests were performed for (F-G), where ****P*<0.001. (H-I) Stromal cells derived from WAT were subjected to adipogenic differentiation in the presence of vehicle, RSPO2 (200 ng/mL), WNT3A (20 ng/mL), or RSPO2 + WNT3A, throughout the duration of differentiation (H) or for the final 24 h of differentiation (I). Oil Red O staining was performed to visualize lipid droplet abundance. Scale bars: 1000 µm. For all qPCR data (C, F-G), gene expression was normalized to *Tbp* housekeeper levels and the vehicle or day 0 condition set to 1, with error bars representing means±SEM.

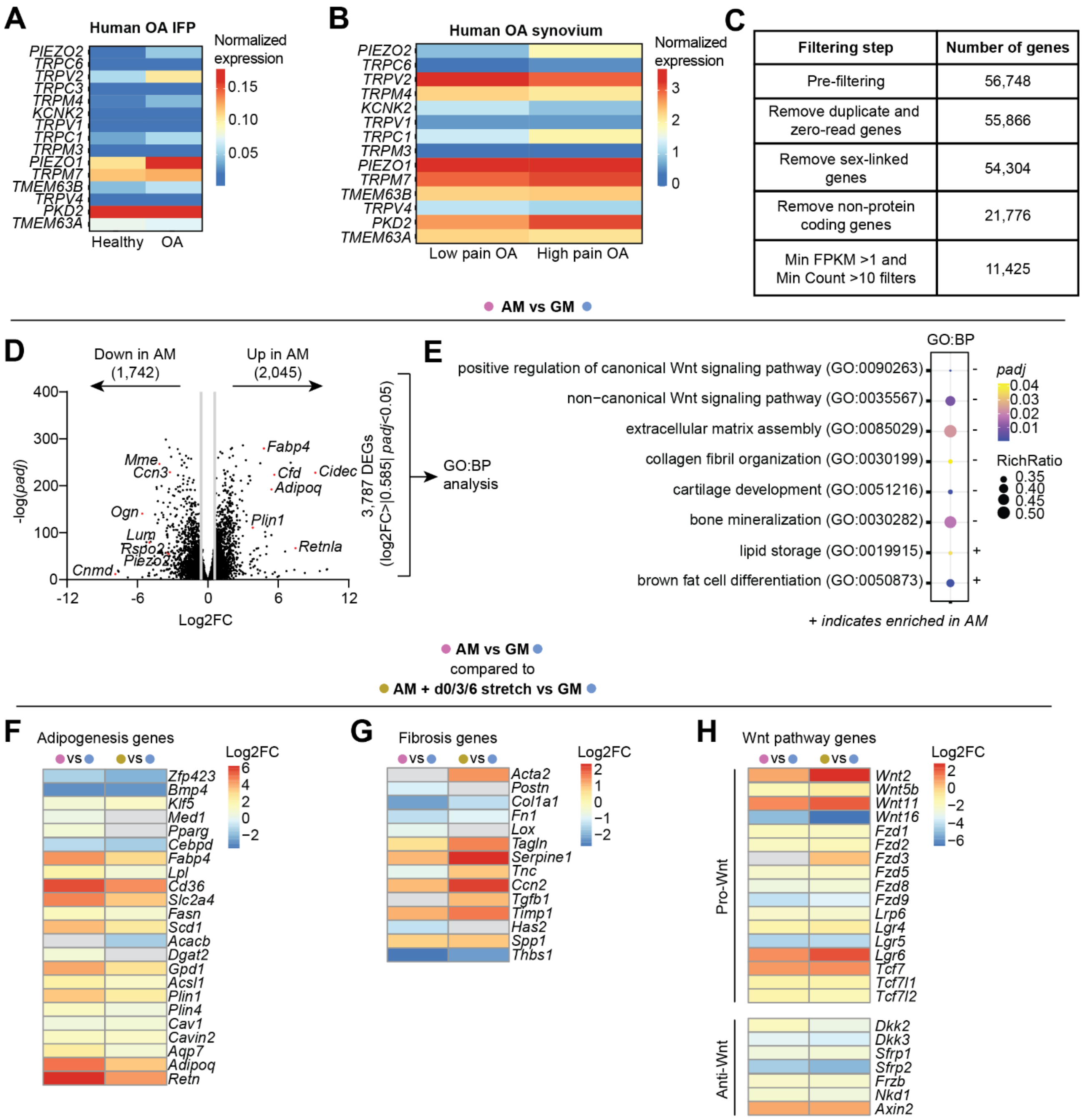


**Fig S5. Mechanical loading blocks adipogenesis. Related to Fig 5.** (A) Log-normalized expression of mechanoreceptor genes in human healthy and OA IFP tissue from pseudo-bulk analysis of publicly available scRNA-seq data, GSE216651. (B) Log-normalized expression of mechanoreceptor genes in human synovium from OA patients reporting low or high knee pain, based pseudo-bulk analysis of publicly available scRNA-seq data, GSE248453. (C) Gene filtering steps and remaining number of genes after each step, for bulk RNA-seq quality control on samples subjected to dynamic loading experiments. (D) Volcano plot showing significant DEGs between AM versus GM (log2FC>|0.585|, *padj*<0.05). Key DEGs are named. (E) Pathway statistical enrichment analysis using the Gene Ontology: Biological Processes (GO:BP) database was performed on the 3,787 significant DEGs from (D). Enriched GO:BP terms are depicted as bubbles, where color represents significance (*padj*) and bubble size indicates RichRatio (DEGs present in the term divided by the total genes in the term). A plus sign (+) indicates enrichment in the AM condition, and vice versa for (-). (F-H) Two-layered differential expression analysis was performed – first, comparing AM to GM, and second, comparing AM + d0/3/6 stretch to GM. Heatmaps were then generated to compare the per-gene log2FC of these two comparisons side-by-side using curated lists of adipogenesis genes (F), fibrosis genes (G), and Wnt pathway genes (H), broken into genes that promote or inhibit Wnt signaling. Color scale represents log2FC, and gray indicates that the gene was not significantly differentially expressed in that comparison.

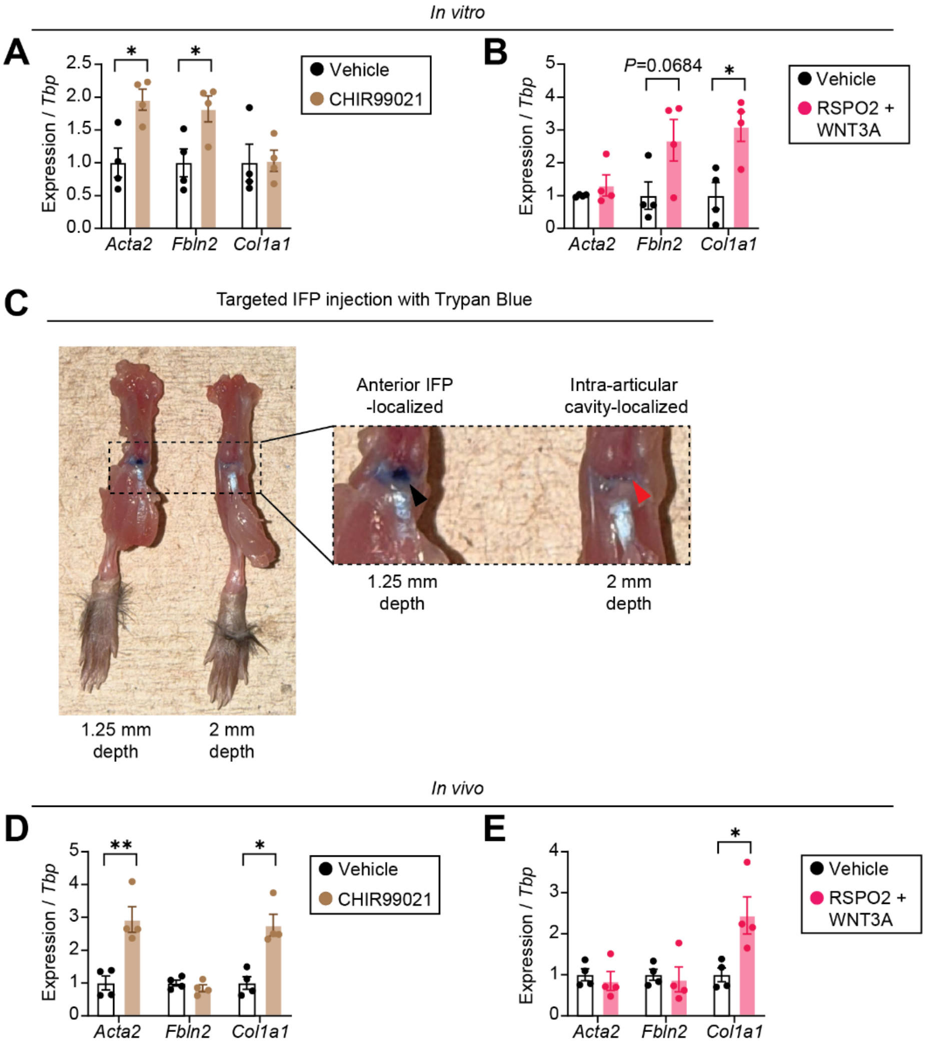


**Fig S6. Chronic Wnt overactivation diminishes intra-articular adipose signature. Related to Fig 6.** (A-B) IFP explants from naïve mice were cultured in adipogenic maintenance medium for 48 h in the presence of 5 µM CHIR99021 or DMSO vehicle (A), or 200 ng/mL RSPO2 + 20 ng/mL WNT3A or PBS vehicle (B) (n=4). Relative expression of fibrotic genes was assessed. (C) Trypan blue was injected intra-articularly at a depth of 1.25 mm or 2 mm. The 1.25 mm injection depth showed preferential accumulation of stain in the IFP (black arrowhead) in the anterior compartment of the joint, whereas the 2 mm depth localized to the intra-articular cavity (red arrowhead). (D-E) Naïve mice received thrice-weekly injections for four weeks, directly into the IFP. 5 µg of CHIR99021 or 5% DMSO/95% PBS vehicle (D), or 500 ng RSPO2 + 50 ng WNT3A or PBS vehicle (E) were delivered in a total volume of 5 µL per injection (n=4). Treatments and their respective vehicles were injected into paired right and left limbs of the same mouse. Relative expression of fibrotic genes was assessed. For (A-B), unpaired two-tailed t-tests were performed, where *$P<0.05$. For (D-E), paired two-tailed t-tests were performed, where *$P<0.05$, **$P<0.01$. For (A-B and D-E), transcript levels were normalized to *Tbp* housekeeper expression, vehicle groups for each gene were set to 1, and error bars are means±SEM.
